# Production of systemically circulating Hedgehog by the intestine couples nutrition to growth and development

**DOI:** 10.1101/002626

**Authors:** Jonathan Rodenfels, Oksana Lavrynenko, Sophie Ayciriex, Julio L. Sampaio, Maria Carvalho, Andrej Shevchenko, Suzanne Eaton

**Author notes:** These authors contributed equally to this work (Oksana Lavrynenko, Sophie Ayciriex, Julio L. Sampaio). present address: Lipotype GmbH, Tanzberg 47, 01307 Dresden, Germany (Julio L. Sampaio). present address: Instituto Gulbenkian de Ciencia, Rua da Quinta Grande, 62780-156 Oeiras, Portugal (Maria Carvalho).

## Abstract

In *Drosophila* larvae, growth and developmental timing are regulated by nutrition in a tightly coordinated fashion. The networks that couple these processes are far from understood. Here, we show that the intestine responds to nutrient availability by regulating production of a circulating lipoprotein-associated form of the signaling protein Hedgehog (Hh). Levels of circulating Hh tune the rates of growth and developmental timing in a coordinated fashion. Circulating Hh signals to the fat body to control larval growth. It regulates developmental timing by controlling ecdysteroid production in the prothoracic gland. Circulating Hh is especially important during starvation, when it is also required for mobilization of fat body triacylglycerol (TAG) stores. Thus, we demonstrate that Hh, previously known only for its local morphogenetic functions, also acts as a lipoprotein-associated endocrine hormone, coordinating the response of multiple tissues to nutrient availability.

## Introduction

Organismal growth and development are regulated by the availability of nutrients. The reaction of a developing animal to nutrient deprivation involves a complex inter-organ response orchestrated by the nervous, endocrine and digestive systems, with the goal to allow basic cellular metabolism but to delay energy demanding processes. A major physiological response to starvation is the utilization of stored energy and the delay of growth and development.

The *Drosophila* fat body, a tissue analogous to mammalian white adipose tissue and liver, has an important role in coupling nutrient availability to growth. In response to changes in the availability of sugars, amino acids and lipids, it releases secreted factors that regulate growth by modulating systemic insulin signaling. (Arquier et al. 2008; Rajan and Perrimon 2012; Geminard et al. 2009; and reviewed in Hietakangas and Cohen 2009). It also acts as a depot for these nutrients, and can release them in response to starvation (reviewed in Telfer and Kunkel 1991; Arrese and Soulages 2010). The fat body directly senses circulating amino acid levels using the amino acid transporter Slimfast and the Target of Rapamycin (TOR) pathway (Colombani et al. 2003). However, how it senses the availability of other types of nutrients is unknown.

In addition to regulating growth, nutrient availability also controls developmental progression by influencing production of ecdysteroids in the prothoracic gland. During larval development, ecdysteroids are produced in pulses that regulate molting and initiate pupariation (reviewed in Thummel 1996). These pulses are tightly coupled to organismal growth. A key developmental decision occurs once larvae achieve a “critical weight” (reviewed in Mirth and Riddiford 2007). After this point, starvation no longer delays pupariation, although it affects the final size of resulting adults. Critical weight is associated with a small increase in ecdysteroid levels, which later rise dramatically to initiate pupariation (Warren et al. 2006). The commitment to and timing of pupariation are subject to multiple checkpoints and can be affected not only by nutrition, but also damage to imaginal tissues, circadian signals, and temperature (reviewed in Shingleton and Mirth 2012). Consistent with this, multiple signaling pathways in the prothoracic gland influence the decision to up-regulate ecdysteroid production (Mirth et al. 2005; Colombani et al. 2005; McBrayer et al. 2007; Rewitz et al. 2009; Gibbens et al. 2011; Cáceres et al. 2011; Ou et al. 2011) How these networks interact to couple ecdysteroids production to the size of the animal is far from understood.

In other organisms, the digestive system is an important node in the signaling network that controls metabolism. Signals originating from the stomach and the intestine activate neural and humoral pathways to regulate food intake and metabolic homeostasis (reviewed in Cummings and Overduin 2007). Less is known about signals produced by the *Drosophila* gastrointestinal tract, although it harbors neuroendocrine cells that can synthesize peptide hormones, and is connected to the nervous system (Cognigni et al. 2011; Miguel-Aliaga 2012; Lemaitre and Miguel-Aliaga 2013). The digestive system is well-placed to directly sense nutrient availability as opposed to circulating nutrients. However, whether the *Drosophila* gastrointestinal tract conveys this type of nutritional information is not known.

Hedgehog (Hh) signaling is known to regulate metabolic processes such as fat storage and glucose uptake in mice (Suh et al. 2006; Pospisilik et al. 2010; Teperino et al. 2012). But the source of the relevant Hh ligand has never been identified. Interestingly, we have shown that circulating lipoproteins in flies and mammals carry not only lipid cargo, but also the lipid-modified signaling molecule Hedgehog (Palm et al. 2013). The tissue source and function of circulating Hh is unknown. In this study, we investigate the source and the function of lipoprotein-associated systemically circulating Hh in *Drosophila*.

## Results

### Midgut derived Hh circulates systemically

To investigate the source of circulating Hh, we characterized the tissue expression patterns of a variety of different Gal4 drivers (suppl. Figure 1A), and used them to knock down Hh production. We then measured Hh levels in the larval hemolymph. Ubiquitous induction of *hh*RNAi using *tubulin*-Gal4 efficiently reduces the level of circulating Hh (suppl. Figure 1B). We next used *en-GAL4, en105*-Gal4 or *hh*-Gal4 to target *hh*RNAi to posterior compartments of larval ectoderm and imaginal discs, which express Hh. These drivers are also active in the salivary glands. Knock-down using these drivers had no effect on the level of circulating Hh, despite strongly reducing *hh* expression in imaginal discs. (Figure 1A, and suppl. Figure 1C). Thus, circulating Hh does not derive from the imaginal discs, larval ectoderm or salivary gland. Therefore, we used immunofluorescence to identify other tissues that might express *hh*. We noted that Hh protein is present in the larval midgut and its attached gastric caeca (Figure 1B). In the midgut, Hh localizes to the enterocytes, which function in nutrient absorption (Figure 1C). To ask whether this region of the gut also transcribes *hh*, we performed *in-situ* hybridization and RT-PCR on dissected midgut samples. These experiments showed that *hh* mRNA is present in the midgut, and that it localizes to enterocytes but not gastric caeca (Figure 1D,E). Thus, Drosophila midgut enterocytes transcribe and translate Hh. The Hh protein found in gastric ceaca must be derived from other sources.

**Figure 1.**
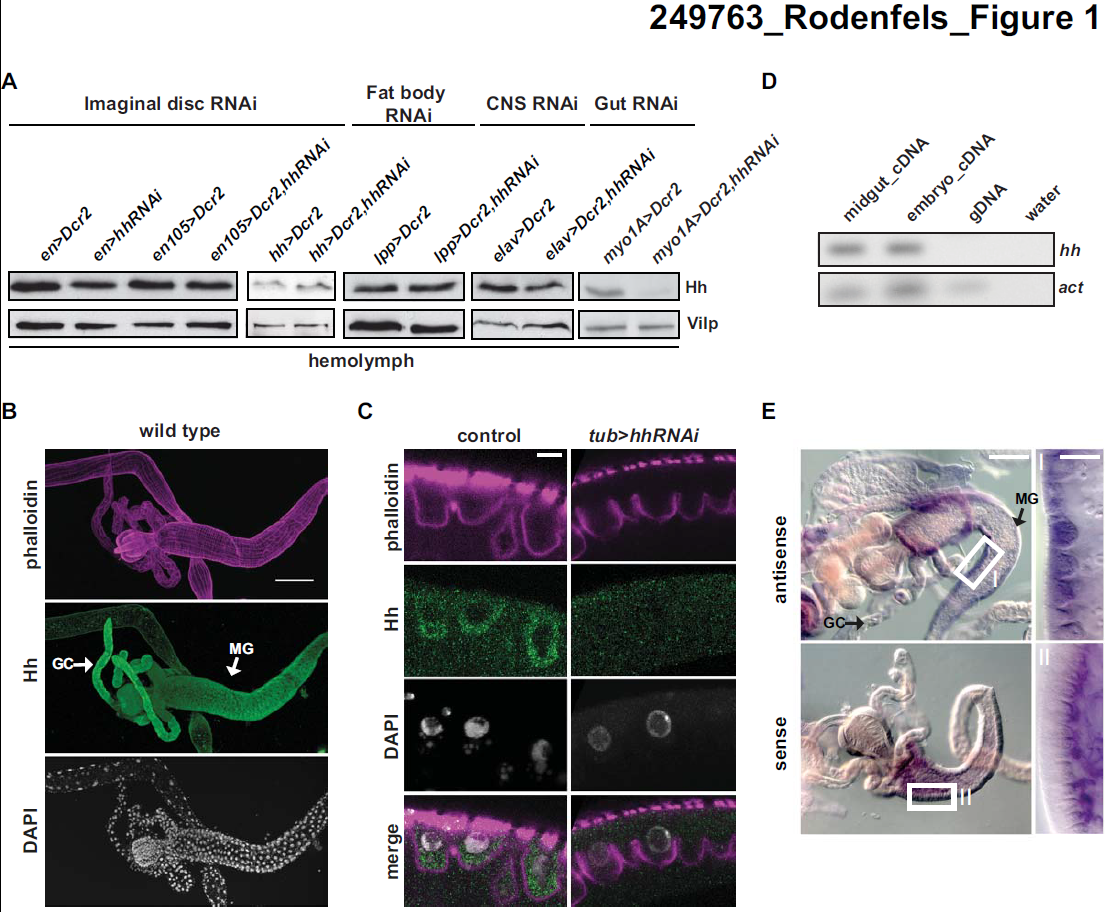
Midgut derived Hh systemically circulates. (**A**) Immunoblot of hemolymph from larvae in which Hh is knocked down with *en* Gal4, *en(105)*-Gal4, *hh*-Gal4, *lpp*-Gal4, *elav*-Gal4, *myo1A*-Gal4). Knock down with *myo1A*-Gal4 strongly reduces Hh levels in the hemolymph. (**B-C**) Immunofluorescence of 2^nd^ instar larval midguts of wt (B,C) and *tub >hh*RNAi (C) animals, stained for Hh F-actin (phalloidin) and DNA (DAPI). GC=gastic ceace, MG=midgut, Scale bars=100 *μ*m (B), 5 *μ*m (C). Ubiquitous knock-down of *hh* with *tub*-Gal4 depletes enterocyte localized Hh. (**D**) RT-PCR for *hh* from second instar midgut samples. Hh message is present in dissected midgut samples. (**E**) *In-situ* hybridization for *hh* mRNA on second instar midgut samples. Higher magnification of the marked areas are displayed on the right (**I+II**) Scale bars= 100 *μ*m and 5 *μ*m marked area. *hh* mRNA localizes to migut enterocytes.

To ask whether circulating Hh originates from the larval midgut, we used the *myo1A*-Gal4 driver. This driver is active in the midgut enterocytes and the salivary gland, and is also expressed in a few cells in the fat body, CNS, and disc peripodial membrane (Figure 1, and suppl. Figure 1A). Driving *hh*RNAi with *myo1A*-Gal4 reduces, but does not completely eliminate, the amount of Hh in the larval hemolymph (Figure 1A). To ask whether the activity of *myo1A*-Gal4 in fat body or CNS underlies its ability to reduce Hh levels in the hemolymph, we induced *hh*RNAi in these tissues more completely using *lpp*-Gal4 or *elav*-Gal4 (which drive strong expression in the fat body and CNS, respectively). Neither reduced systemic Hh levels when used to drive *hh*RNAi (Figure 1A). These results strongly suggest that midgut enterocytes are a major source of systemic Hh, although minor amounts may come from other tissues.

### Circulating Hh inhibits systemic growth and slows developmental timing

Although larvae lacking circulating Hh can give rise to normally patterned adults, these animals are somewhat smaller than controls. We therefore wondered whether Hh might influence larval growth or developmental timing. To investigate this, we first examined the larval growth rate of animals homozygous for a temperature-sensitive mutation of *hh* (*hh^ts2^*). At the restrictive temperature, *hh^ts2^* animals grow faster and pupariate earlier than two different wild type controls (suppl. Figure 2A). They pupariate just as efficiently as wild type animals (supple. Figure S2D), and faster growth and earlier pupariation combine to produce *hh^ts2^* pupae of normal size (although *hh^ts2^* animals do not give rise to viable adults) (suppl. Figure 2A). Thus, in addition to its other patterning functions in the larva, Hh acts to slow the rate of larval growth and delay pupariation. To ask whether loss of circulating Hh was responsible for these effects, we asked whether the speedier growth and development of *hh^ts2^* larvae could be reversed by expression of Hh in midgut enterocytes. Indeed, driving *hh* expression using *npcGal4* in the *hh^ts2^* mutant background completely reverses the accelerated pupariation and growth of *hh^ts2^* larvae (Figure 2A). In fact, these animals pupariate even later than wild type controls. Pupal volume is indistinguishable from that of wild type in all cases, suggesting that delayed pupariation is accompanied by a reduction in the larval growth rate (Figure 2B). Thus, *hh^ts2^* mutant larvae develop more rapidly than wild type because they lack Hh expression in the midgut.

**Figure 2.**
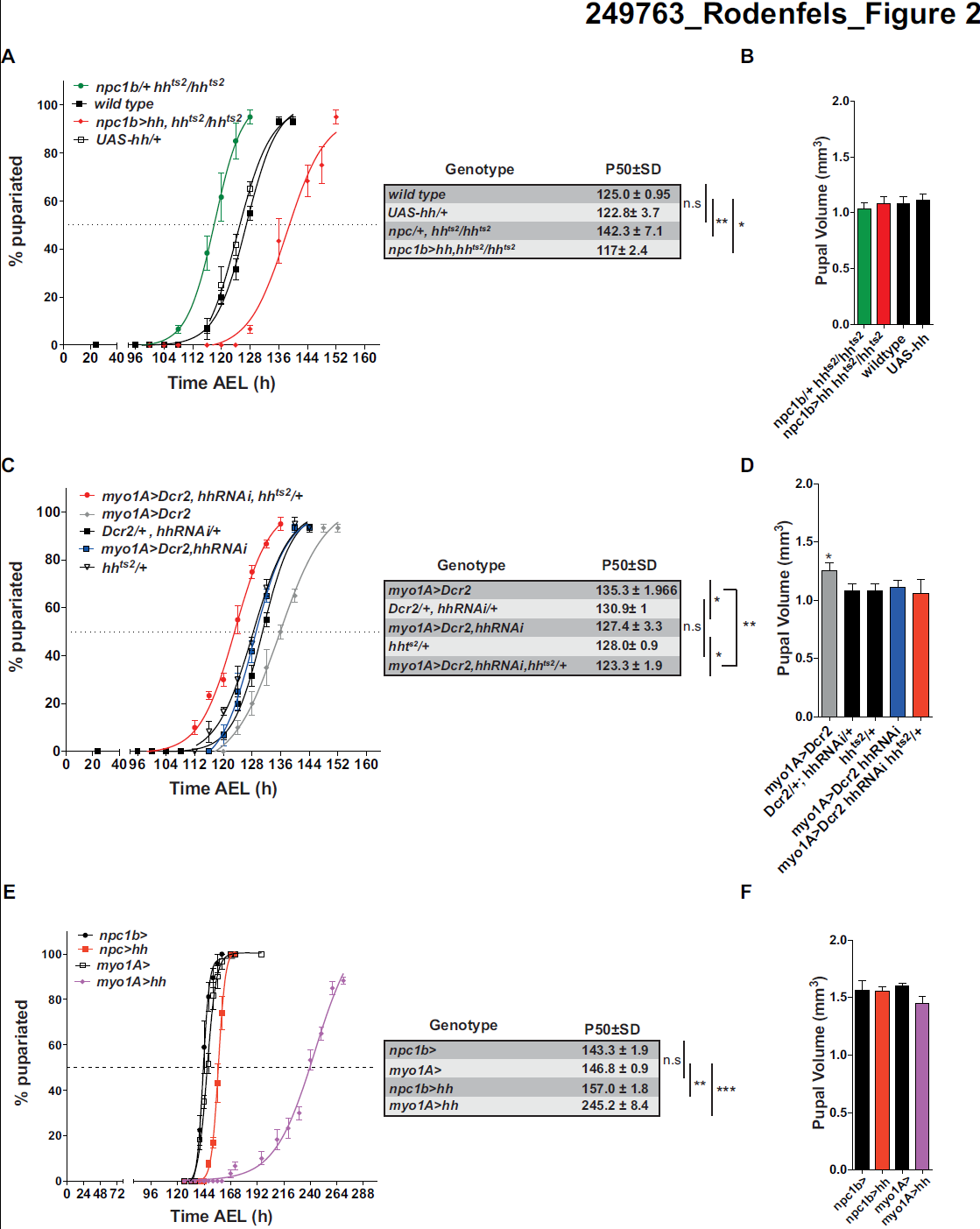
Circulating Hh inhibits systemic growth and slows developmental timing. (**A,C,E**) Percentage of pupariated animals at different times (in hours) after egglaying (AEL). The time AEL at which 50% of larvae have pupariated (P50) was determined by non-linear regression for each biological replicate. P-values are based on students t-test comparing genotype specific P50 values. Error bars represent standard deviation, n=3 *p<0.05, **p<0.01, ***p<0.001 (**B,D,F**) Measurement of pupal volume of animals with increased or decreased circulating Hh levels. Error bars represent standard deviation, n=3, *p<0.05

To ask whether reducing Hh production in the midgut influenced developmental timing, we used *myo1A-Gal4* to drive *hhRNAi* in midgut enterocytes, and enhanced the efficiency of RNAi by co-expressing *Dicer2 (Dcr2)*. This treatment reduces but does not eliminate circulating Hh. These animals pupariate significantly earlier than animals expressing *Dcr2* alone (Figure 2C, suppl. Figure 2B). However *myoGAL4 >Dcr2* larvae develop more slowly than other control larvae. Thus, it was difficult to be sure that reduced Hh production in midgut enterocytes had increased the rate of larval development. To investigate this further, we asked whether removing one wild type *hh* allele could further accelerate the development of *myoGAL4 >Dcr2, hhRNAi* animals. Indeed, RNAi-mediated *hh* knock-down in a *hh^ts2^*/+ heterozygous background caused significantly earlier pupariation than knockdown in a wild type background. (Figure 2C). Larvae pupariate at the same volume (Figure 2D), suggesting that the growth rate is also increased by performing Hh knock-down in a hh^ts2^/+ heterozygous background. Taken together, these results suggest that lowering production of Hh in the midgut speeds the rate of larval growth and pupariation.

To ask whether elevating midgut Hh production could slow larval development of otherwise wild type animals, we expressed *hh* under the control of two different Gal4 drivers: *myo1A-Gal4* and *npc1b-Gal4. Npc1b*-Gal4-driven expression is strictly midgut specific and strongly increases the level of circulating Hh. *Myo1A-Gal4* is expressed more highly than *npc1b*-Gal4 in the midgut, but is also active a few other tissues. *Myo1A-Gal4* driven *hh* expression increases circulating Hh levels more dramatically than *npc1b*-Gal4 (Palm et al. 2013). Pupariation of *npc1b >hh* larvae is delayed by 14 hours, compared to control *npc1b*/+ larvae (Figure 2E). Myo1A-driven *hh* has a stronger effect, delaying pupariation by 4 days compared to controls (Figure 2E). Delayed pupariation in *hh* over-expressing animals is accompanied by a proportional reduction in larval growth, such that these animals pupariate at close to a normal size (Figure 2F, suppl. Figure 2C).

Taken together, these data show that the amount of Hh circulating in the hemolymph controls both the larval growth rate and the speed of pupariation in a proportional fashion.

### Circulating Hh promotes survival under starvation

Growth and developmental timing are modulated in response to nutrient availability. Circulating Hh is produced in the tissue where nutrient absorption occurs. Therefore, we investigated whether its production responded to nutrient uptake. Quantitative RT-PCR and Western blot analysis show that midgut *hh* expression is significantly elevated upon starvation in both pre and post-critical weight larvae (Figure 3A,B). To distinguish the effects of protein, sugar and lipids on *hh* expression, we asked which nutrients could reverse the upregulation of *hh* expression in starving larvae. Adding 5% sugar (fructose or glucose) to starvation medium (PBS/Agar) did not reverse *hh* upregulation (Figure 3C). Adding 10% chloroform-extracted yeast autolysate as a source of amino acids, and other water-soluble molecules, partially reversed *hh* upregulation (Figure 3C). However, adding both sugar and protein consistently and completely reversed Hh upregulation in the midgut (Figure 3C). These data suggest that midgut cells deprived either of sugar or of amino acids increase *hh* expression. Interestingly, dietary lipids do not appear to be required to keep *hh* expression low in the midgut.

**Figure 3.**
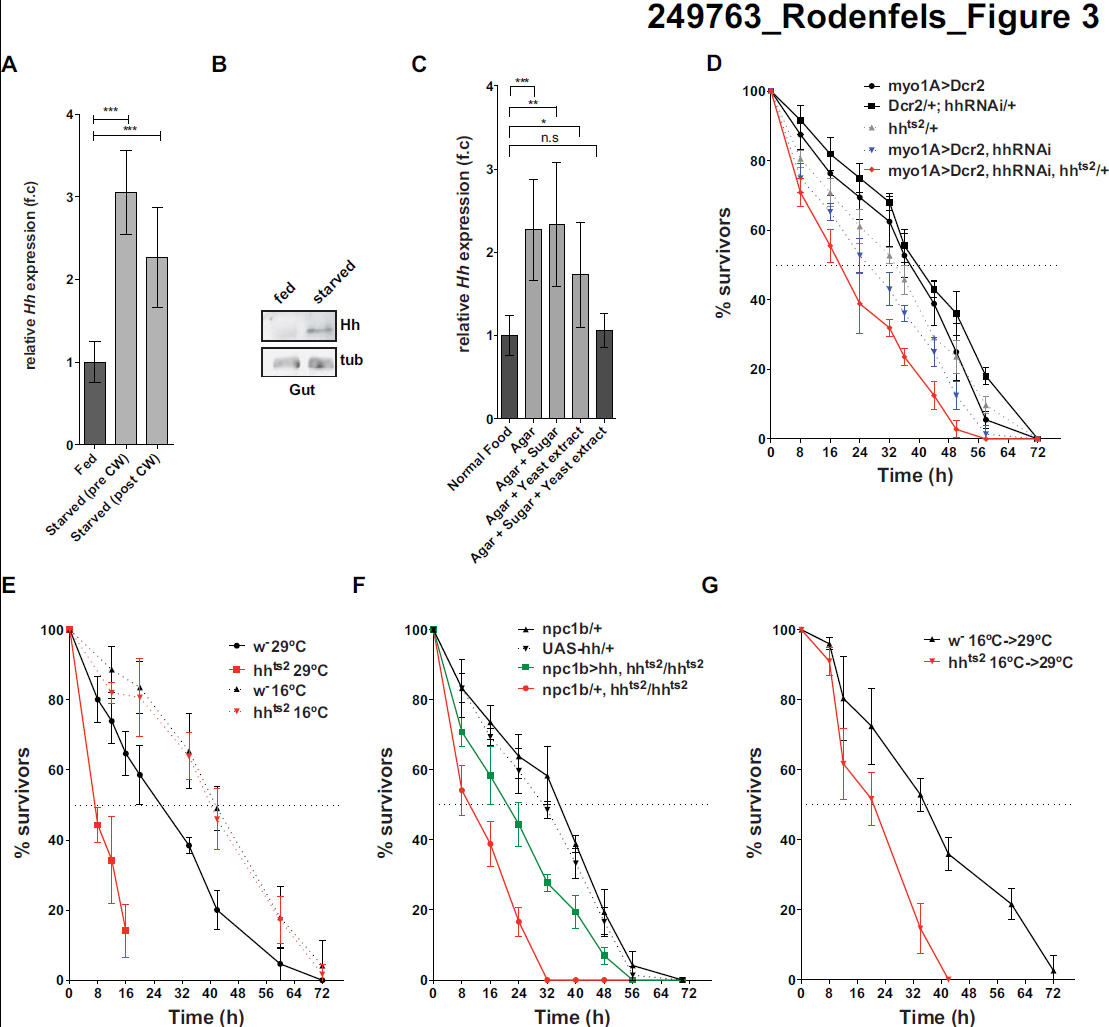
Midgut Hh promotes survival under starvation conditions. (**A**) Measurements of relative midgut *hh* mRNA expression by qRT-PCR of animals that were either fed or starved (pre-and post-critical weight). *Hh* expression was normalized to *rp49*. Fold change are relative to fed controls. n>15, *p<0.05, **p<0.01, ***p<0.001. (**B**) Immunoblot for Hh of midguts of fed or starved. Hh becomes enriched in starved midguts. (**C**) Measurements of relative midgut *hh* mRNA expression by qRT-PCR of animals that were either fed, starved, or fed sugar, chloroform extracted yeast autolysate or sugar plus chloroform extracted yeast autolysate. *Hh* expression was normalized to *rp49.* Fold changes are relative to fed controls. n=6, *p<0.05, **p<0.01, ***p<0.001. (**D**) Survival rate upon starvation of *myo1A>Dcr2,Dcr2/+,hhRNAi/+,hh^ts2^/+ myo1A>Dcr2,hhRNAi* and *myo1A>Dcr2,hhRNAi,hh^ts2^/+late* 2^nd^ instar larvae cultured at 29°C. (**E**) Survival rate upon starvation of wild type and *hh^ts2^* late 2^nd^ instar larvae cultured at permissive (16°C) or at restrictive temperature (29°C) after embryonic development until time of starvation. (F) Survival rate upon starvation of control (*npc1b/+, UAS-hh/+*) and *npc1b/+, hh^ts2^/hh^ts2^* and *npc1b>hh, hh^ts2^/hh^ts2^* cultured at restrictive temperature (29°C) (**G**) Survival rate upon starvation of wild type and *hh^ts2^* cultured at permissive (16°C) until the late 2^nd^ instar followed by starvation at the restrictive temperature (29°C).

Control larvae can survive for 40 hours when they are starved before reaching critical weight (Figure 3D). To ask whether Hh production by the midgut was important for survival under starvation conditions, we reduced Hh function in larvae starved before critical weight by driving *hh*RNAi in the midgut with *myo1A*-Gal4. Although *myo1A*-driven *hh*RNAi does not reduce survival of normally fed animals (suppl. Fig 2E), it reduces survival on starvation from 40 to 24 hours (Figure 3D).. To confirm that starvation sensitivity was caused by reduced Hh production in the midgut, we asked whether the phenotype could be enhanced in a *hh^ts2^*/+ heterozygous background. Indeed, removing one functional copy of *hh* decreased survival even further (Figure 3D) Thus, even the incomplete reduction of circulating Hh caused by *myolA*-induced *hh*RNAi strongly decreases survival upon starvation. This suggests that circulating Hh promotes survival under starvation conditions.

To reduce systemic Hh levels more completely, we blocked Hh function using the temperature sensitive Hh allele *hh^ts2^. Hh^ts2^* larvae cultured at the restrictive temperature (29°C) from hatching onwards complete larval development with no increased lethality when fed normally, but die during early pupal stages (suppl. Fig 2D). Thus, Hh function is not required for larval survival or pupariation of normally fed animals. However, when starved before critical weight, these animals die 38 hours earlier than wild type controls (Figure 3E). This reduction in viability is due to the temperature sensitive mutation because *hh^ts2^* larvae do not differ from wild type controls when starved at the permissive temperature (16°C) (Figure 3E).

To ask whether starvation sensitivity of *hh^ts2^* homozygous larvae was due to lack of Hh production in the midgut, we asked whether midgut-specific Hh expression could rescue survival of starved *hh^ts2^* homozygous animals. Indeed, midgut-specific Hh expression significantly improves survival on starvation. (Figure 3F). Thus, midgut Hh expression increases survival in starving larvae.

We wondered whether circulating Hh acted at the time of starvation to increase larval survival. Alternatively, loss of circulating Hh during larval growth might affect nutrient storage, making them more sensitive to later starvation. To distinguish these possibilities, we shifted *hh^ts2^* larvae to 29°C coincident with starvation, and monitored their survival. These animals die 24 hours earlier than wild type (Figure 3G). Taken together, these results show that increased production of circulating Hh by the midgut upon starvation is essential for normal survival under starvation conditions.

### Hh produced in the midgut is not required for midgut homeostasis or midgut tracheal branching

We wondered whether Hh produced in the midgut was required locally to influence intestinal function. To ask whether altered Hh levels in the midgut might affect its structure or homeostasis, we assayed morphology, proliferation and apoptosis in the midgut. We also counted the number of undifferentiated adult midgut precursor (AMPs) cells within a cluster, and total AMPs clusters in a given region of interest. Neither *hh* overexpression nor RNAi-mediated knock-down of *hh* in the midgut induced apoptosis or changed enterocyte cell size or morphology (suppl. Figure 3A). Furthermore, the total number of AMP clusters and the number of within a given cluster remained constant (suppl. Figure 3A,B), consistent with previous results (Jiang and Edgar 2009). Thus, Hh produced by midgut enterocytes does not locally influence their morphology or homeostasis.

To more directly assess whether Hh signaling might affect the function of ECs or AMPs, we autonomously increased or decreased pathway activity in these cell types by expressing RNAi or dominant negative/active constructs for Hh pathway components under the control of *myo1A*-Gal4 and *esg*-Gal4, respectively. The Hedgehog receptor, Patched (Ptc), represses signaling when Hh is not present. Binding of Hh to Ptc blocks Ptc-mediated repression of the 7-pass transmembrane protein Smoothened (Smo), which can mediate both transcriptional and non-transcriptional pathway outputs (van den Heuvel and Ingham 1996; Alcedo et al. 1996; Chen et al. 2001; DeCamp et al. 2000; Zhang et al. 2001; Ogden et al. 2008). We either activated Hh signaling using *ptc*RNAi, or inhibited it by *smo*RNAi in ECs or in AMPs. None of these manipulations altered cell morphology, apoptosis or proliferation in the midgut (suppl. Figure 3C,D). To further assess whether Hh signaling in ECs and AMPs could regulate systemic growth and developmental timing we measured the timing of pupariation and the resulting pupal volume after increasing or decreasing Hh signaling activity in midgut EC and AMPs. None of these manipulations affected pupal volume or the timing of pupariation (suppl. Figure 3E,F).

Taken together, these results strongly suggest that Hh produced in the midgut is not required locally in either ECs or AMPs to regulate systemic growth and developmental timing. We wondered whether Hh signaling in the midgut might influence the ability of larvae to cope with starvation. To test this, we either activated or inhibited Hh signaling in enterocytes and monitored survival upon starvation. Neither inhibition nor activation of Hh signaling in enterocytes affects starvation sensitivity (suppl. Fig 3G).

Recently, the regulated branching of midgut trachea has been implicated in the regulation of energy stores and their mobilization in response to nutrients.. (Linneweber et al. 2014). To test whether midgut Hh expression might regulate starvation sensitivity by affecting the structure of midgut trachea, we investigated their morphology in larvae with altered Hh expression in the midgut. Examining trachea in 3 different regions of the midgut revealed no effect of *hh* overexpression or knock-down on tracheal size or morphology (suppl. Figure 4A). Furthermore activation or inhibition of hh signaling in trachea terminal cells did not affect survival upon starvation (suppl. Figure 4B). Thus, Hh produced by midgut enterocytes does not appear to influence the overlying trachea.

### Circulating Hh does not influence local wing imaginal disc Hh signaling

Is circulating Hh required for normal Hh signaling in the imaginal discs? The growth of imaginal discs exerts strong non-autonomous effects on larval developmental timing (Garelli et al. 2012; Colombani et al. 2012). Changes in imaginal disc Hh signaling due to circulating Hh might thereby indirectly affect larval developmental timing. We ask whether endogenously circulating Hh was required for normal patterning in the wing disc, we reduced circulating Hh levels in the gut using *myo1A*-Gal4 and monitored Hh levels and pathway activity in the wing disc. Hh is locally expressed in the posterior compartment of the wing disc, and spreads into the anterior compartment where its receptor *ptc* is expressed. Hh signaling stabilizes the full-length form of the Cubitus interruptus (Ci) transcription factor (Ci^155^), and upregulates transcription of target genes including *ptc* (Tabata and Kornberg 1994; Aza-Blanc et al. 1997). Reducing Hh levels in the hemolymph did not affect the distribution of Hh in the wing disc (Figure 4A,B). Neither did it change the range of Ci^155^ stabilization or *ptc* expression (Figure 4A,B), and flies emerged with normally patterned wings (not shown). Thus, endogenous levels of circulating Hh are not required for local patterning in wing imaginal discs. While we have previously observed that strong elevation of circulating Hh levels perturbs local patterning in the wing imaginal disc (Palm et al. 2013), more moderate increases produced by expressing Hh under the control of *npc1b*-Gal4 have no effect (Figure 4A,B). Nevertheless, *npc1b*-driven *hh* expression reduces larval growth and delays pupariation (Figure 2C). Thus, circulating Hh does not act indirectly through imaginal discs.

**Figure 4:**
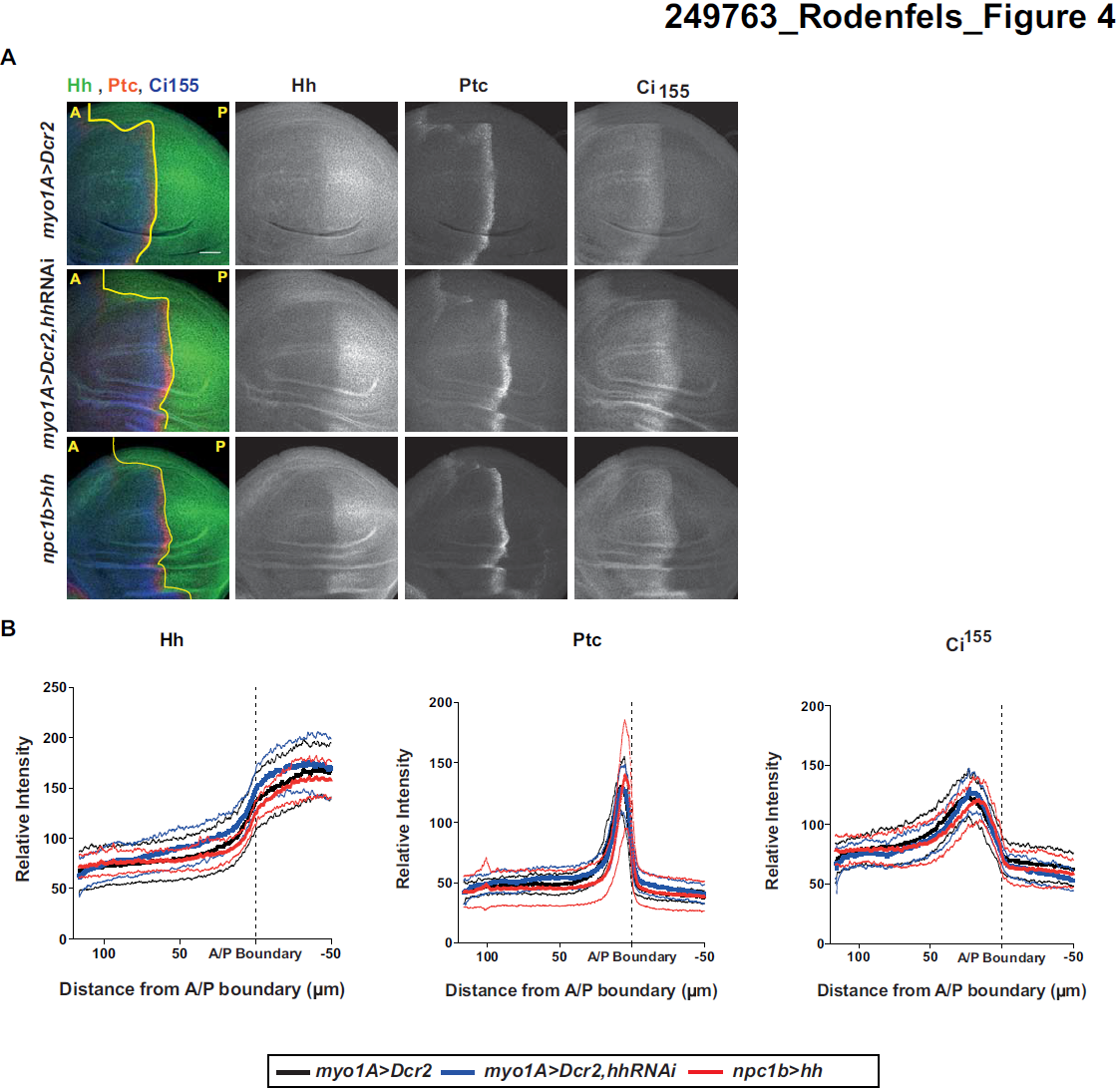
Circulating Hh does not influence local wing imaginal disc Hh signaling. (**A**) Immunofluorescence of 3^rd^ instar wing disc stained for Hh, Ptc and Ci^155^ from *myo1A>Dcr2, myo1A>Dcr2,hhRNAi* and *npc1b>hh* animals. Scale bar=30 *μ*m. Anterior (A)/posterior (P) compartment boundary is marked with a yellow line. (B-D) Quantification of Hh (B), Ptc (C) and Ci^155^ (D) levels of control (*myo1A>Dcr2*, black line) animals and animals with decreased (*myo1A>Dcr2, hh-RNAi*, blue line) and increased (*npc1b>hh*, red line) midgut Hh levels, n=13, dashed lines represent standard deviation.

### Circulating Hh does not influence the frequency of mouth hook contractions

The mammalian intestine produces hormones that act on the brain to regulate feeding behavior (Sandoval et al. 2008; Strader and Woods 2005).To ask whether circulating Hh produced by the midgut might influence larval feeding, we counted larval mouth hook contractions. Neither overexpression nor RNAi-mediated knockdown of *hh* in the midgut changed the frequency of larval mouth hook contractions (suppl. Fig 4C). Thus, changes in growth and developmental timing are not a simple consequence of less enthusiastic feeding.

### Circulating Hedgehog promotes neutral lipid mobilization in the fat body upon starvation

Since Hh signaling can reduce TAG levels in both flies and mammals, we wondered whether the influence of circulating Hh on starvation resistance, growth rate or pupariation timing was mediated through effects on fat metabolism. We first examined how loss or gain of circulating Hedgehog influenced TAG levels in normally fed animals, using shotgun mass spectrometry. We observed no change in the amount of stored TAG under either condition – either in the fat body or in whole animals (Figure 5 A,B). Thus, altering the levels of circulating Hh is insufficient to change TAG metabolism in well-fed animals. These observations suggest that Hh influences growth and developmental timing by mechanisms that do not depend on TAG metabolism.

**Figure 5.**
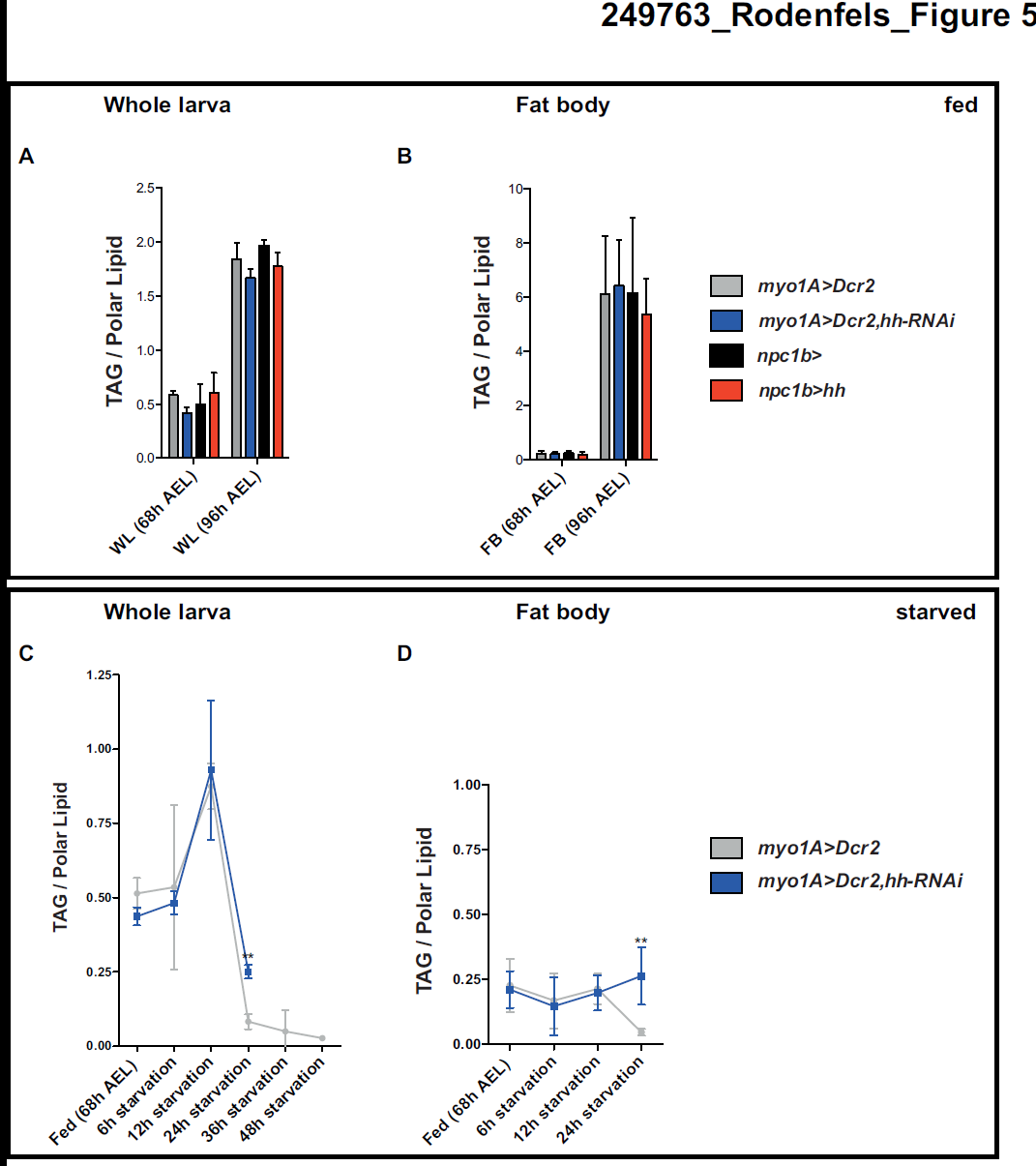
Circulating Hedgehog promotes neutral lipid mobilization in the fat body upon starvation. (**A-B**) Whole animals (A) and fat body (B) TAG levels of normal fed late 2^nd^ instar (68h AEL) and 3^rd^ instar (96h AEL) larvae with reduced or increased Hh levels in the midgut, quantified by mass spectrometry (n=3). TAG levels were normalized to the total amount of membrane lipids. AEL. (**C-D**) Changes in whole animals (C) and fat body (D) TAG levels upon starvation of late 2^nd^ instar larvae with reduced Hh levels in the midgut, quantified by mass spectrometry (n=3). *p>0.05, **p>0.01, ***p>0.001. Error bars represent SD

We next investigated whether circulating Hh might influence TAG metabolism under starvation conditions. In control larvae, starvation initially increases levels of whole larval TAG within 12 hours by almost 2-fold (Figure 5C). Interestingly, TAG’s do not increase in the fat body at this time - this increase must occur in other tissues (Figure 5D). Over the next 12 hours, both whole body and fat body TAG’s drop dramatically (Figure 5C,D). Wild type larvae normally die about 24 hours after TAG stores have dropped (48 hours after starvation) (Figure 4D).

Reducing the levels of circulating Hh during starvation completely blocks mobilization of fat body TAG’s (Figure 5D). However it does not prevent the disappearance of TAG’s from other tissues; whole body TAG’s drop between 12-24 hours to levels almost as low as those in control animals (Figure 5C). Residual whole body TAG at this time likely reflects failed mobilization of fat body stores. These larvae die 24 hours earlier than control larvae – right after the time that wild type animals would have mobilized fat body TAG stores. Thus, circulating Hh is required to mobilize fat body TAG stores under starvation conditions. However elevating circulating Hh in well-fed animals is not sufficient by itself to reduce TAG’s in the fat body – other signals active during starvation must work together with Hh to accomplish this.

### Fat body Hh signaling regulates starvation resistance and larval growth

We wondered whether some of the effects of circulating Hh might be mediated by direct signaling to the fat body – indeed, the fat body expresses Hh pathway components, but not *hh* itself (Suh et al. 2006; Palm et al. 2013; Pospisilik et al. 2010 and suppl. Fig 5A). To investigate this, we first asked whether Hh protein was detectable in the fat body, despite the fact that it is not expressed there. Immunoblotting shows that the fat bodies of feeding third instar larvae indeed contain small amounts of Hh protein (Figure 6A). Fat body Hh protein levels, but not *hh* transcription, increase upon starvation (Figure 6A, and suppl. Figure 5B). These are precisely the conditions under which Hh production is upregulated in the midgut (Figure 3A,B). To determine whether Hh in the fat body was derived from the midgut, we reduced Hh production there and monitored Hh protein in the fat body of starving larvae by Western blotting. Indeed, *hh* knock-down in the midgut reduces Hh protein levels in the fat body (Figure 6B). Thus, the midgut is the source of Hh ligand found endogenously in the fat body. Furthermore, driving *hh* over-expression in the gut using *npc1b*-Gal4 elevates the amount of Hh in the fat body (Figure 6A). Thus, the amount of Hh accumulating in the fat body directly reflects the amount of Hh produced by the midgut.

To ask whether Hh signaling in the fat body might mediate effects on growth, pupariation and starvation resistance, we autonomously increased or decreased pathway activity in this tissue by expressing RNAi or dominant negative/active constructs for Hh pathway components under the control of *lpp*-Gal4. We first examined whether Hh signaling in the fat body controlled TAG metabolism or starvation resistance. Although manipulation of the pathway at the level of the systemic Hh ligand does not alter TAG stores, autonomous pathway activation and inactivation in the fat body cause opposing changes in whole animal TAG – consistent with previous reports (Suh et al. 2006; Pospisilik et al. 2010 and suppl. Fig 5C). Interestingly, both pathway activation and pathway repression reduced survival upon starvation (Figure 6C,D). These animals may die for opposite reasons. When Hh signaling is permanently activated in the fat body, larvae have less stored TAG to mobilize. When it is inactivated, they may be unable to mobilize it.

**Figure 6.**
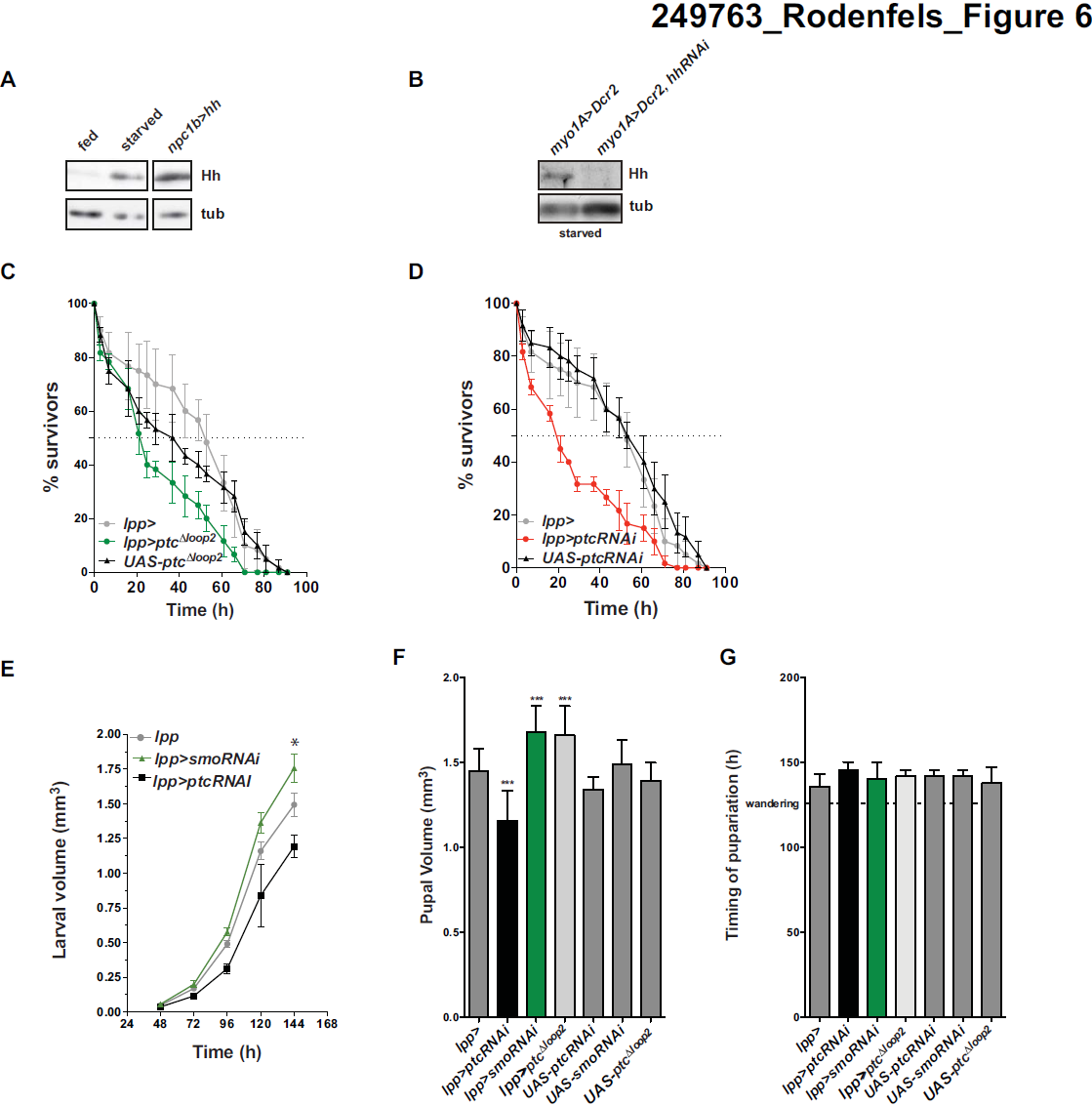
Fat body Hh signaling regulates starvation resistance and larval growth. (**A**) Immunoblot for early 3^rd^ instar fat body Hh from animals that have been normally fed, starved or with increased *hh* expression in the midgut. (**B**) Immunoblot for fat body Hh upon starvation from animals with reduced midgut Hh expression. (**C-D**) Survival rate upon starvation of *lpp>ptc*^Δ*loop2*^ (B) and *lpp>ptcRNAi* (C) and their respective control late 2^nd^ instar larvae. (**E**) Changes in larval volume and determination of the timing of pupariation of animals with increased (lpp>ptcRNAi) or decreased (*lpp>smoRNAi*) Hh signaling specifically in the fat body. Asterix marks white pupal stage and volume. (**F**) Measurement of pupal volume of animals with increased or decrease Hh pathway activity in the fat body. (**F**) Measurements of timing of pupariation of animals with increased or decrease Hh pathway activity in the fat body.

Is the fat body responsible for the alterations in growth and/or developmental timing caused by perturbing *hh* expression in the midgut? To test this, we monitored the timing of pupariation and resulting pupal volume after increasing or decreasing Hh signaling in the fat body. Activating the pathway by RNAi-mediated *ptc* knock-own decreased the larval growth rate (Figure 6E), but did not affect pupariation timing (Figure 6E,G). As a consequence, pupal volume was decreased (Figure 6E,F). Pathway inhibition caused by *smo*RNAi increased larval growth rate (Figure 6E), again without affecting the timing of pupariation (Figure 6E,G), resulting in larger pupae (Figure 6F). Similar results were obtained when Hh signaling was blocked by expression of the constitutively active *ptc*^Δ*loop2*^ (Figure 6F,G). Interestingly, these effects of Ptc and Smo in the fat body appear to be exerted by a non-canonical pathway that does not involve processing of the transcription factor Cubitus Interruptus (Ci) (suppl. Text 1 and suppl. Figure 6). Instead, up and down-regulation of Hh causes opposing changes in specific fat body diacylglycerol species (suppl. Figure 6H). Taken together, these data show that the fat body mediates the effects of circulating Hh on larval growth, and does so by a non-canonical pathway. However, its effects on developmental timing must be exerted through a different tissue.

### Hh signaling in the prothoracic gland regulates developmental timing

The timing of Ecdysteroid production by the prothoracic gland regulates developmental transitions including pupariation. We therefore asked whether Hh might signal to the prothoracic gland to control developmental timing. We first investigated whether Hh pathway components were present in this tissue using immunofluorescence. We detected Ptc, Smo and Ci^155^ specifically in the prothoracic gland, but not in the corpus allatum or corpus cardiacum, which produce other hormones (Figure 7A,B). Furthermore, Lpp, which carries circulating Hh, also accumulates specifically on cells of the prothoracic gland (Figure 7A,B). However Hh itself could not be detected by immunofluorescence (data not shown). To ask whether Hh produced in the midgut was able to travel to the ring gland, we expressed either *hh-gfp* or membrane tethered *cd8-hh-gfp* under the control of *npc1b*-Gal4 and stained ring glands with an antibody to GFP. Hh-GFP is clearly detected on cells of the prothoracic gland, but not when the midgut expresses comparable levels of the membrane-tethered version (Figure 7C, suppl. Fig 7A). Thus, Hh made in the midgut can travel to the prothoracic gland, which expresses Hh pathway components.

**Figure 7.**
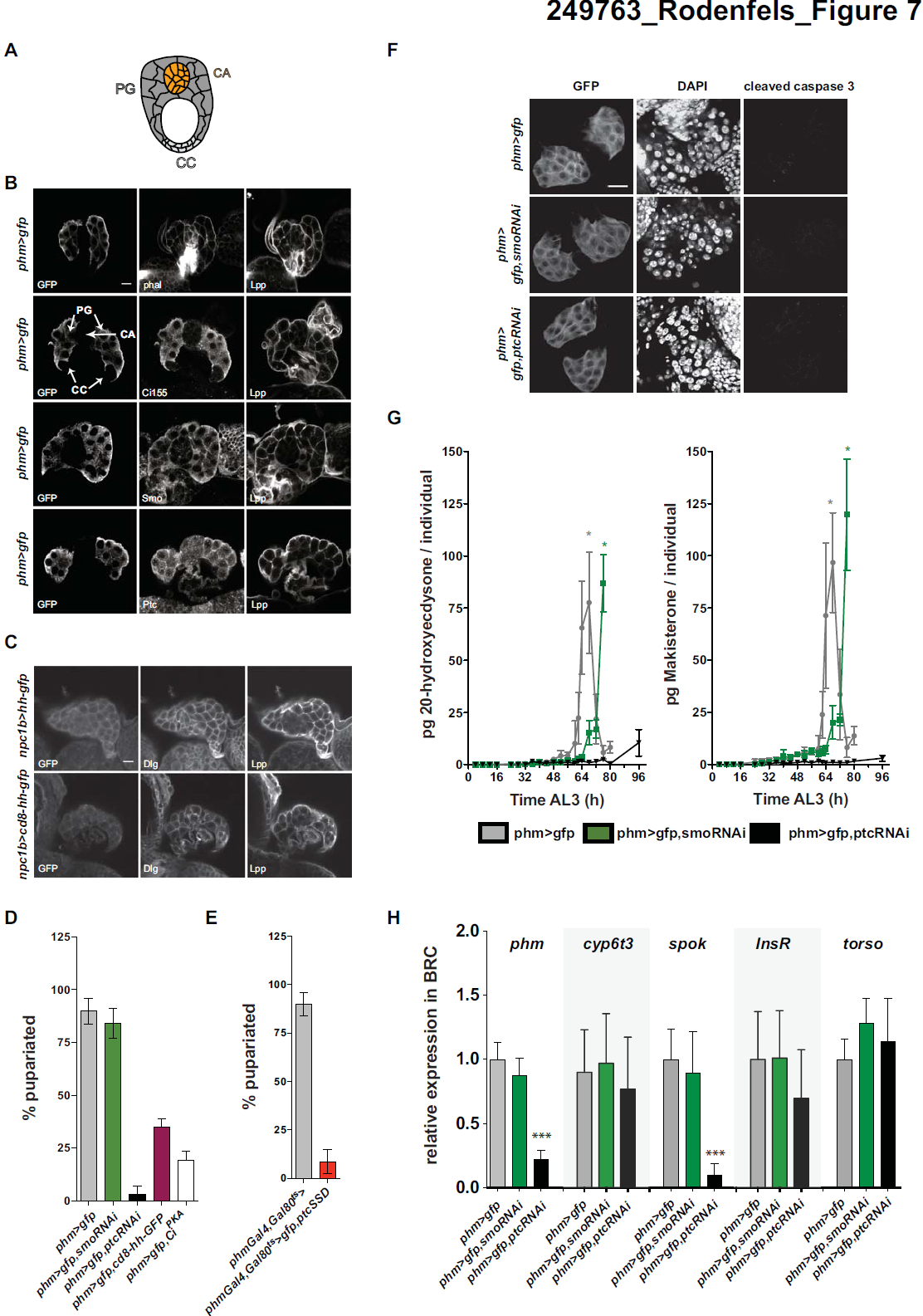
Prothoracic gland Hh signaling regulates ecdysteroid production. (**A**) Cartoon of a *Drosophila* ring gland. Prothoracic gland (PG), corpus cardiac (CC), corpus allatum (CA). (**B**) Immunofluorescence of *phm>CD8-GFP* 2^nd^ instar ring gland stained for GFP, phalloidin, Ci155, Smo, Ptc and Lpp. Scale bar=20μm. (**C**) Immunofluorescence of *npc1b>cd8-hh-gfp* and *npc1B>hh-gfp* 2^nd^ instar ring gland stained for GFP, Discs large (Dlg) and Lpp (**D**) Percentage of larvae pupariating with inhibited (*phm>gJp,smoRNAi*) or activated (*phm>gfp,ptcRNAi*, phm>gfp, cd8-hh-gfp, phm>gfp,Ci^PKA^) Hh signaling pathway in the PG. Error bars represent SD. (**E**) Percentage of larvae pupariating with activated (*phmGal4,Gal80^s^>gfp,ptc^SSD^*) Hh signaling pathway in the PG. Error bars represent SD. (**F**) Immunofluorescence of phm>gfp, *phm>gfp,smoRNAi* and *phm>ptcRNAi* 2^nd^ instar ring gland stained for GFP, DNA (DAPI) and cleaved caspase 3, Scale bar=25*μ*m. (**G**) Quantitative ecdysteroid MS time course for 20E (**E**) and Maki A from whole larva with inhibited (*phm>gfp,smoRNAi*) or activated (*phm>gfp,ptc*RNAi) Hh signaling pathway in the PG or from control animals (*phm>gfp*) asterisk marks white pupal stage, Error bars indicate SD of biological triplicates (**H**) Quantitative RT-PCR for *phm, cyp6t3, spok, InsR and toros* at 96h AEL from dissected brain-ring gland complexes (BRC) with increased or decreased Hh pathway activity in the PG relative to control (*phm>gfp*) BRCs. Error bars represent SD of biological triplicates *p< 0.05, **p< 0.01 and ***p< 0.001.

To investigate the autonomous effects of Hh signaling in the prothoracic gland, we used *phm*-Gal4 to drive the expression of constructs that activate or inactivate Hh signaling. We then monitored the rate of larval growth, timing of pupariation, and starvation sensitivity. We used four methods to activate Hh signaling: RNAi-mediated knock-down of *ptc*; expression of the dominant negative, *ptc^SSD^* ; expression of membrane-tethered Hh; and expression of dominant active Ci, Ci^PKA^. Each of these treatments inhibits pupariation, and loss of Ptc function blocks it almost completely (Figure 7D,E). In contrast, inhibiting Hh signaling by *smo*RNAi expression only slightly delays puparation (8-12 hours after wild type controls) and larvae pupariate at the same size (suppl. Figure 7B). These data suggest that loss of Hh signaling alone is not sufficient to induce early pupariation, but that increased Hh signaling in the prothoracic gland inhibits pupariation. In contrast to its function in the fat body, Hh signaling in the prothoracic gland does not strongly influence larval growth or starvation sensitivity (suppl. Figure 7C,D).

Taken together these results suggest that the physiological effects of elevating circulating Hh can be accounted for by its combined action in the fat body and prothoracic gland – Hh signaling in the fat body reduces larval growth, and Hh signaling in the prothoracic gland delays pupariation.

How does Hh signaling in the prothoracic gland block pupariation?

Expressing *ptcRNAi* or *smoRNAi* in the prothoracic gland does not disturb its morphology or cause Caspase activation, thus cell death in the prothoracic gland cannot account for the effects of *ptcRNAi* on pupariation (Fig 7F). *PtcRNAi* reduces nuclear area by 20%, suggesting that these cells may be slightly less polyploid than those in wild type prothoracic glands (suppl. Figure 7E).

Pupariation is normally initiated by a rise in the level of ecdysteroid hormones. At early wandering stages, the prothoracic gland increases transcription of ecdysteroid biosynthetic enzymes and elevates production of ecdysone, which is hydroxylated by other tissues to 20-OH ecdysone. To ask whether Hh signaling in the prothoracic gland blocked pupariation by preventing production of ecdysteroids, we quantified ecdysteroids over the course of the third instar and in early pupae using LC-MS/MS. In wild type third instar larvae, we detect both 20-OH ecdysone and the related hormone, Makisterone A, but not ecdysone or methyl-ecdysone (the precursor of Makisterone A) (Figure 7G). This suggests that 20-hydroxylation must be rapid and efficient at this stage. 20-OH ecdysone and Makisterone A levels begin to rise 56 hours after the molt from L2 to L3, and peak in white prepupae (Figure 7G). In contrast, larvae with constitutively active Hh signaling in the prothoracic gland fail to produce high levels of either 20-OH ecdysone or Makisterone A (Figure 7G). Only a slight rise in hormone levels is apparent even 96 hours after the molt from L2 to L3, long after wild type larvae have pupariated. Thus, constitutive Hh signaling in the prothoracic gland blocks 20-OH ecdysone and Makisterone A production. Reducing Hh signaling by *smo* knock-down in the PG only slightly delays pupariation (suppl. Figure 7B). Consistent with this, these animals increase production of 20-OH ecdysone and MakisteroneA with slightly delayed kinetics compared to wild type (Figure 7G).

Quantitative RT-PCR shows that expression of ptcRNAi and ptcSSD in the prothoracic gland reduces mRNA levels for the ecdysteroid biosynthetic enzymes *phantom (phm) and spookier (spok)*, (Warren et al. 2004) (Figure 7H, suppl. Figure 7F). However, expression of cyp6t3 a potential rate-limiting enzyme in ecdysteroid biosynthesis remains unchanged (Figure 7H, suppl. Figure 7F). This suggests that reduced transcription of biosynthetic enzymes contributes to lower ecdysone/methyl-ecdysone biosynthesis in the prothoracic gland.

Both Prothoracicotropic hormone (PTTH) and Insulin signaling can regulate the onset of Ecdysone production (Mirth et al. 2005; Rewitz et al. 2009). To ask whether Hh signaling might disturb either of these processes, we quantified expression of Insulin receptor (*InsR*) or the Ptth receptor *torso* by qRT-PCR in brain-ring gland complexes. Neither *ptcRNAi* nor expression of *ptc^SSD^* significantly changed the expression of *insR* or *torso,* suggesting that Hh signaling does not act by reducing susceptibility to PTTH or *Drosophila* Insulin-like peptides (Figure 7H, suppl. Figure 7F. Thus, high Hh signaling reduces transcription of ecdysteroid biosynthetic enzymes and inhibits production of Ecdysone and Makisterone A, delaying pupariation.

Taken together, these data suggest that the increase in ecdysteroid biosynthesis that precedes and initiates pupariation does not occur if Hh signaling in the prothoracic gland is too high. However, reducing Hh signaling is not sufficient to initiate ecdysteroid biosynthesis. Lowered Hh signaling is likely only one of many inputs that regulate this important decision.

## Discussion

The ability to couple nutrition to growth and developmental progression allows animals to thrive in unpredictable environments. Work in *Drosophila* has begun to outline how different organs communicate to coordinate these processes. Key interactions have been demonstrated between the fat body, prothoracic gland and central nervous system (Mirth et al. 2005; Colombani et al. 2005; Geminard et al. 2009; Delanoue et al. 2010). Our results highlight the importance of the intestine as another node in the signaling network. The intestine is well-placed to specifically sense intake of nutrients as opposed to their circulating levels, which may reflect other physiological conditions and not food availability alone.

We have shown that Hh pathway activation in the prothoracic gland inhibits ecdysone production and delays pupariation – consistent with the delay in pupariation caused by increasing circulating Hh, In contrast, although reducing circulating Hh causes larvae to pupariate sooner, inhibiting Hh pathway activation in the prothoracic gland is not sufficient to do so. Clearly, the information directly provided to the prothoracic gland by lower circulating Hh levels must be integrated with that of other signals. For example, Insulin, TGFβ and PTTH signaling are all required to increase ecdysteroid production (Mirth et al. 2005; Colombani et al. 2005; McBrayer et al. 2007; Rewitz et al. 2009; Gibbens et al. 2011). It seems likely that the drop in circulating Hh is sensed by other tissues, which influence the production of such confirmatory signals.

Circulating Hh also signals to the fat body to slow larval growth, and to mobilize TAG’s during nutrient deprivation. Its action in the fat body and prothoracic gland are independent of each other. Growth and pupariation timing can be altered separately by autonomously activating Hedgehog signaling in either of these tissues by itself. Thus, circulating Hh couples systemic growth and developmental timing by its independent actions in the fat body and the prothoracic gland. While Hh acts directly on both these tissues, other studies have clearly shown that information can also flow back and forth between the ring gland, the fat body, and Dilp-producing neurons in the brain. Humoral signals such as Activin, and the fat body-derived Leptin homologue Unpaired are transmitted to the brain and promote the release of Dilp’s. These increase insulin signaling in the ring gland, which elevates ecdysteroid production (Rajan and Perrimon 2012; Geminard et al. 2009);(Mirth et al. 2005) (Ghosh and O'Connor 2014; Gibbens et al. 2011). Conversely, moderately increased levels of ecdysteroids act on the fat body to inhibit myc activity, which slows larval growth – possibly by reducing Dilp release (Colombani et al. 2005; Delanoue et al. 2010). Taken together, the signals flowing between the gut, fat body, IPC’s and prothoracic gland represent a robust information processing network that produces sensible decisions about growth and development.

We have highlighted the importance of intestinal derived circulating Hedgehog to regulate lipid mobilization and to couple systemic growth to developmental progression during nutritional stress. Interestingly, work by others shows that Hh transcription is increased by bacterial infection in the gut, and also by loss DHR96 (Chakrabarti et al. 2012; Bujold et al. 2010). This nuclear hormone receptor mediates the response to xeniobiotics and regulates TAG metabolism (King-Jones et al. 2006; Sieber and Thummel 2009). Thus it would be interesting to test whether circulating Hh might regulate growth and development in response to other physiological stresses.

A variety of different Hh secretion forms have been reported(Zeng et al. 2001; Chen et al. 2004; Tanaka et al. 2005; Panáková et al. 2005; Creanga et al. 2012; Ohlig et al. 2012; Palm et al. 2013). We have previously studied two different secretion forms – one that is sterol-modified and lipoprotein associated, and another that is monomeric and not sterol modified (Palm et al. 2013). These different forms exert different and synergistic effects on the signaling pathway in *Drosophila* imaginal discs (Palm et al. 2013). In contrast, signaling in the fat body is mediated entirely by the Lpp-associated form. In disc, the Lpp form of Hh blocks the repressive activity of lipoprotein lipids on Smoothened stability, but even high levels do not influence target gene activation by Cubitus Interruptus – this requires the non-sterol modified form (Palm et al. 2013). Consistently, the action of lipoprotein-associated Hedgehog in the fat body requires Ptc and Smo, but does not appear to act through Ci-dependent transcription. These findings emphasize the physiological importance of the lipoprotein-associated Hh form, and suggest that the fat body will be a good system to further study its unique effects on the Hh pathway. It will be interesting to know whether Lpp-associated Hh might directly influence lipid storage, or regulate the availability of signaling lipids in the fat body, as suggested by work in imaginal discs. (Khaliullina et al. 2009; Palm et al. 2013).

Recently, the a non-canonical Hedgehog signaling pathway has been implicated in regulating lipid and sugar metabolism in adult animals – including mice and flies (Pospisilik et al. 2010; Teperino et al. 2012). However, whether this occurs in response to a Hedgehog ligand, and what the source of such a ligand might be, was completely unknown. Here we show that circulating *Drosophila* Hh, produced by the intestine, regulates lipid and steroid hormone metabolism and adapts the animal to its nutritional environment. Since the presence of Hedgehog in circulation is conserved (Palm et al. 2013), it may be that circulating mammalian Shh exerts a similar function. It will be particularly interesting to also examine the impact of circulating Shh on adrenal steroidogenesis, given its function in *Drosophila* and the dramatic effects of glucocorticoids on lipid and sugar metabolism (Walker 2007). Interestingly, the production of Hh proteins in the gut is widely conserved – even in Cnidarians (van den Brink 2007; Matus et al. 2008). Thus, it seems possible that a metabolic function for circulating and lipoprotein-associated Hedgehog proteins might have arisen early in evolution.

## Material & Methods

### Fly Stocks

The following stocks were used: *lpp*-Gal4 (Brankatschk and Eaton 2010), *en(105)*-Gal4, *en*-Gal4 (Eugster et al. 2007), *hh*-Gal4, *tub*-Gal4, *myo1A*-Gal4 (Jiang et al. 2009), *npc1b*-Gal4 (Voght et al. 2007), *phm*-Gal4 (Ono et al. 2006), *hh^ts2^*, UAS-*hh* (Strigini and Cohen 1997), UAS-*ptc*^Δ*loop2*^ (Briscoe et al. 2001),*UAS-ptc^SSD^ (gift from I. Guerrero)*, UAS-Ci^PKA^ (Méthot and Basler 1999) drsf-Gal4 (Gervais and Casanova 2011), *esg*-Gal4, *UAS-cd8-gfp (this study)*, wild-type Oregon R (OreR) and *w^−^* were obtained from Bloomington. UAS-*Dicer2, smoRNAi, ptcRNAi* were obtained from VDRC, Vienna, and *hh*RNAi was obtained from DGRC, Kyoto.

### Survival upon starvation

To measure survival, larvae from 4h egg collections were kept on normal food until 60h after egg laying (AEL). For experiments with *hh^ts2^* animals, eggs were collected at 22°C for 4h and were kept at permissive temperature (16°C) for 48h to allow embryonic development. Hatched *hh^ts2^* and control larvae were shifted to restrictive temperature either at 48h AEL or just prior to starvation. 2^nd^ instar larvae were washed in PBS and transferred to 24 well plates (a single larva per well) filled with Agar/PBS 0.8% (w/v). Dead larvae were counted every 8h and the percentage of surviving animals from individual 24 well plates was plotted over time. For each genotype, three 24 well plates were quantified.

### Growth rates and developmental timing analysis

Growth rate and developmental timing analysis was performed as described (Delanoue et al. 2010; Colombani et al. 2005). In brief, larvae of different genotypes from a 4h egg collection were synchronized at hatching (23h AEL), cultured in petri dishes (6cm) filled with normal food at a density of 50 larvae per dish. Digital images were captured every 24h and the larval or pupal volume was estimated by quantifying the area of the length and the width of the animals in Fiji, then estimating the corresponding volume using the formula 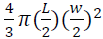 (L=larval length, w= larval width). The average volume of a least 15 animals was plotted over time in biological triplicates. Timing of pupariation was defined as the time at which 50% of the larvae have pupariated.

### Lipids extraction and shotgun mass spectrometry

Lipids were extracted from fat body and from whole larvae and quantified by shotgun mass spectrometry as described in (Carvalho et al. 2012) with modifications. Briefly, samples were homogenized in 150 mM ammonium bicarbonate buffer. From each homogenate, a volume containing ~ 2 nmol of lipid material was removed and extracted using a modified Folch protocol at 4°C (WL n=3, FB n=4 fed to 12h of starvation, n=6 for 24h of starvation experiment).

Mass spectrometry analyses were performed on a Q Exactive instrument (Thermo Fisher Scientific, Bremen, Germany) equipped with a robotic nanoflow ion source TriVersa NanoMate (Advion BioSciences,Ithaca, NY) using chips with the diameter of spraying nozzles of 4.1 mm. The ion source was controlled by Chipsoft 8.1 software. Ionization voltage was set at +0.95 kV and −0.95 kV and back pressure was set at 1.25 and 0.6 psi in positive and negative modes, respectively. The temperature of ion transfer capillary was 200°C; tube voltages were 90 V (MS^+^) and −150V (MS^-^). Acquisitions were performed at the mass resolution *R_m/z400_*=140 000. AGC control was set at 10^6^ ions and maximum injection time to 250 ms.

Dried lipid extracts were re-suspended in 100 μL chloroform-methanol mixture 1:2 (v/v). For the analysis, 10 μL of samples were loaded onto 96-well plate (Eppendorf, Hamburg) and diluted either with 13 μL of 13.3 mM ammonium acetate in 2-propanol for positive ion mode analysis or with 10 μL of 0.01% methylamine in methanol for negative ion mode analysis. The plate was sealed with aluminum foil. Each sample was analyzed for ca 1 min in positive and negative ion modes: PE, PC, PC-O, TAG, CerPE and DAG were quantified in positive and PI, PS, PG, PE, PEO-, Cer, HexCer in negative mode.

Lipids were identified by LipidXplorer (Herzog et al. 2011) software by matching *m/z* of their monoisotopic peaks to the corresponding elemental composition constraints. Molecular Fragmentation Query Language (MFQL) queries compiled for all the aforementioned lipid classes are available at the LipidXplorer wiki site: https://wiki.mpicbg.de/wiki/lipidx/index.php/Main_Page. Mass tolerance was 5 ppm and intensity threshold was set according to the noise level reported by Xcalibur software (Thermo Scientific).

The content of glycerolipids was normalized per polar lipids and expressed in pmol lipid x/pmol polar lipids. Lipids standards were purchased from Avanti Polar Lipids (Alabaster, Alabama). Solvents were purchased from Sigma-Aldrich (Taufkirchen, Germany).

### Quantification of ecdysteroids by LC-MS/MS

Larvae of different genotypes from a 4h egg collection were synchronized at hatching (23h AEL), cultured in petri dishes filled with normal food at a density of 50 larvae each plate. Larvae were staged again at the 2nd to 3rd instar transition (AEL3). Every 4h groups of 4 animals were collected in 100 μl H_2_O and smashed with plastic pestle and extracted overnight with 1 ml of cold (6°C) methanol. Samples were centrifuged and supernatant was collected. The pellet was twice re-extracted with 1 ml methanol for 1 hour, the extracts combined, cleaned by solid-phase extraction, dried down and analyzed by LC-MS/MS as described in (Hikiba et al. 2013) with minor modifications. The detection limit of the method was *ca* 5 pg per individual ecdysteroid. Lipids standards were purchased from Avanti Polar Lipids (Alabaster, Alabama). Solvents were purchased from Sigma-Aldrich (Taufkirchen, Germany).

For additional Material & Methods see supplemental material

## Acknowledgments

We are grateful to Bruce Edgar, Michael O’Connor, Isabell Guerrero and the Bloomington Stock Center for fly stocks. We gratefully acknowledge Robert Holmgren and the Developmental Studies Hybridoma Bank for providing antibodies. We thank Sven Ssykor and Marko Brankatschk for help with transgenesis. We thank Andreas Sagner, Marko Brankatschk and Jochen Rink for critical comments on the manuscript.

## Supplemental Material

Rodenfels et al., Production of systemically circulating Hedgehog by the intestine couples nutrition to growth and development

### Itemized List of Supplemental Material

Suppl. Legend Fig.1: Provides information about expression pattern of Gal4 lines used in Fig 1., and shows controls of Fig 1.
Suppl. Legend Fig. 2.: Provides additional growth and developmental timing data in support of Fig. 2. Shows that knock-down of *hh* with *myo1A-Gal4* does not affect viability of normal fed animals, and does not affect viability of *hh^ts2^* larvae until pupal stage
Suppl. Legend Fig.3: Control experiment to show that Hh produced in the midgut does not act on local midgut enterocytes and adult midgut progenitors cells to regulate systemic growth and developmental timing and starvation sensitivity
Suppl. Legend Fig.4: Control experiment to show that Hh produced in the midgut does not regulate tracheal terminal cell morphology and feeding behavior.
Suppl. Legend Fig.5: Shows controls for Fig. 6
Suppl. Legend Fig.6: Results to show that the fat body mediates the effect of circulating Hedgehog on larval growth by a non-canonical Hh pathway.
Suppl. Legend Fig.7: Shows control experiments for Fig. 7 and provides additional qRT-PCR data for ptcSSD expressed in the PG.
Suppl. Material and Methods: Antibodies, Hemolymph preparation, Fly husbandry and fly food, Generation of *UAS-hh-gfp* and *UAS-cd8-Hh-gfp*, Immunohistochemistry of larval tissues, *In-situ* hybridization, RNA extraction and reverse transcription polymerase chain reaction, Quantitive RT–PCR, Quantification of mouth-hook contractions, Quantification of midgut adult midgut progenitor cell (AMP) clusters.
Suppl. References: References within suppl. Material

**Supplemental Figure 1.**
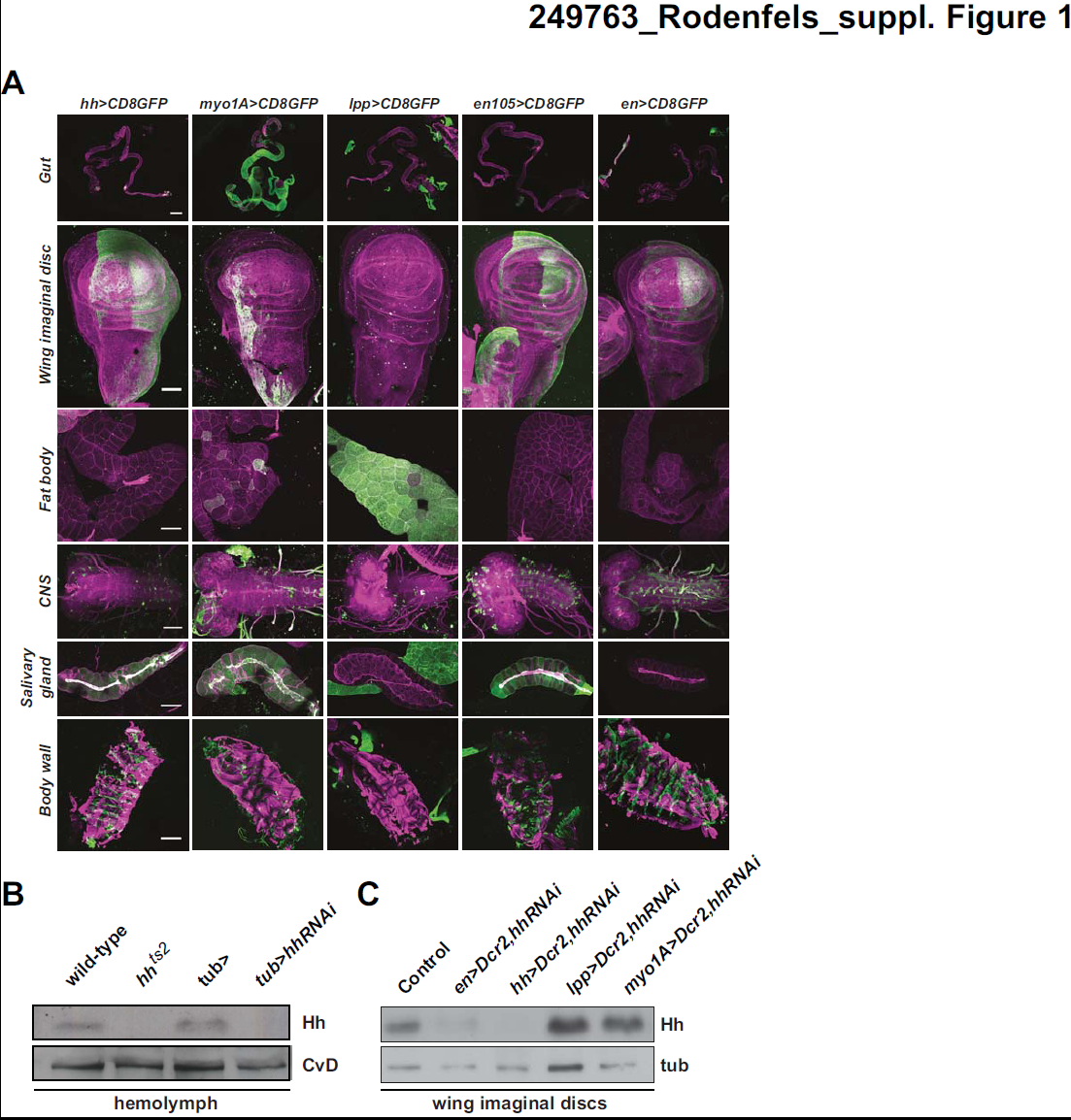
(**A**) Immunofluorescence for GFP of 2^nd^ instar midguts (scale bar=300 *μ*m), 3^rd^ instar wing imaginal discs (scale bar=50 *μ*m), 3^rd^ instar fat body (scale bar=100 *μ*m), 3^rd^ instar central nervous systems (scale bar=50 *μ*m), 3^rd^ instar salivary glands (scale bar=200 *μ*m) and 2^nd^ instar body walls (scale bar=200 *μ*m) of animals expression *uas-cd8-gfp* under the control of *hh*-Gal4, *myo1A*-Gal4, *lpp*-Gal4, *en(105)*-Gal4 and *en*-Gal4 (**B**). Immunoblot for Hh of 3^rd^ instar hemolymph samples of wt, Hh^ts2^ cultured at 29C after embryonic development, *tub>, tub>hh*RNAi animals. Hh is present in wt and *tub>* hemloymph. Hh is depleted in Hh^ts2^ and *tub>hh*RNAi animals (**C**) Immunoblot for Hh of 3^rd^ instar wing imaginal discs of wt, *en>Dcr2,hh*RNAi, *hh> Dcr2,hh*RNAi, *lpp>Dcr2,hh*RNAi and *myo1A>Dcr2,hh*RNAi animals. *En*-Gal4 and *hh*-Gal4 driven *Dcr2,hh*RNAi efficiently depletes wing imaginal disc Hh. *Lpp*-Gal4 and *myo1A*-Gal4 driven *Dcr2,hh*RNAi does not alter Hh levels in wing discs.

**Supplemental Figure 2.**
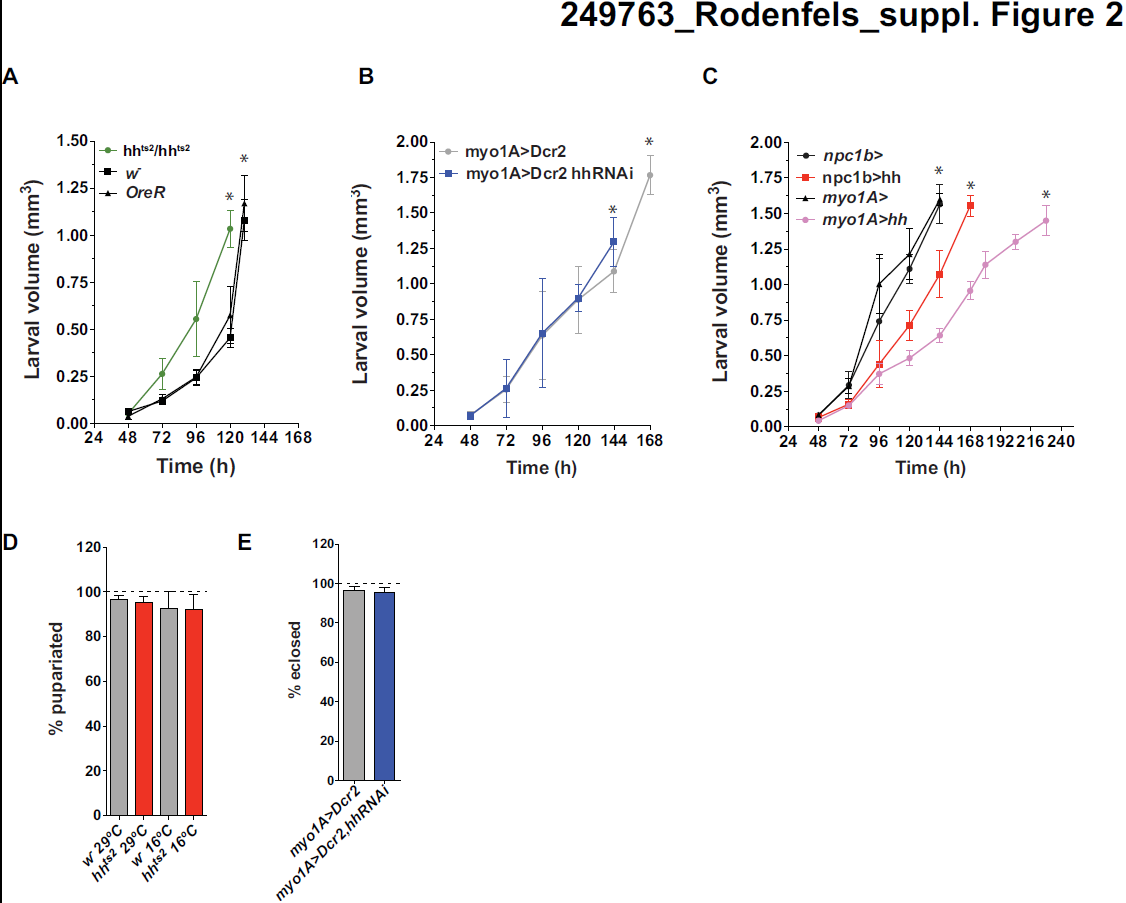
(**A-C**) Changes in larval volume and determination of the timing of pupariation of wild-type (*w^−^* and *OreR*) and *hh^ts2^* animals cultured at the restrictive temperature (29°C) (A) and of animals with reduced (*myo1A>Dcr2,hhRNAi*) (B) or increased (*npc1B>hh* and *myo1A>hh*) (C) *hh* expression in the midgut with their respective control animals. Asterix marks white pupal stage and volume. (**D**) Percentage of normally fed wild type and *hh^ts2^* larvae that pupariate when cultured at permissive (16°C) or restrictive temperature (29°C). Loss of Hh does not affect the percent of larvae that pupariate under normal feeding conditions. (**E**) Percentage of of *myo1A>Dcr2* and *myo1A>Dcr2,HhRNAi* animals that produce adults when normally fed. *Hh* knock down in the gut does not affect viability under normal feeding conditions.

**Supplemental Figure 3.**
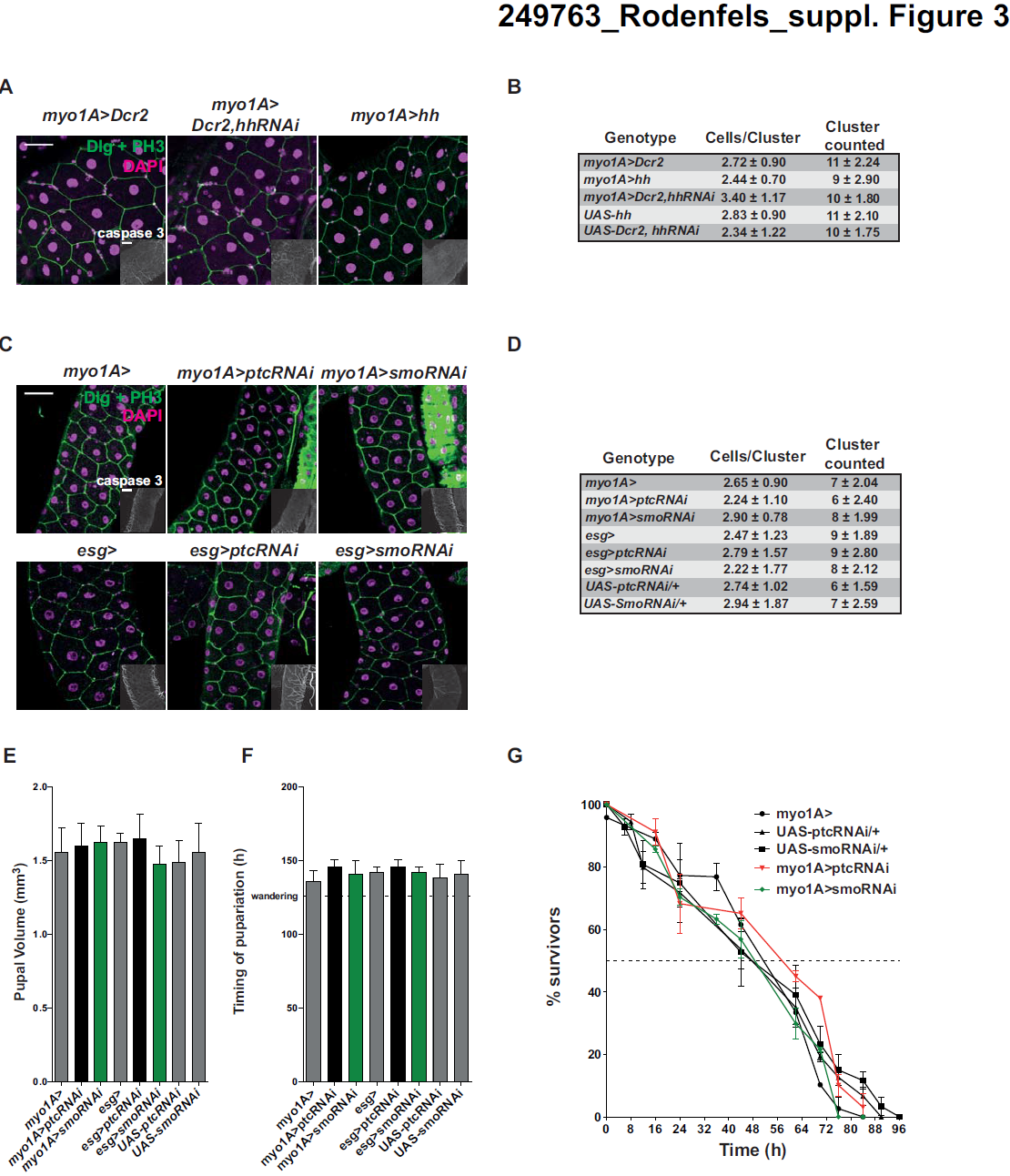
(**A**) Immunofluorescence of 3^nd^ instar larval midguts of *myo1A>Dcr2, myo1A>Dcr2,HhRNAi*, and *myo1A>hh* animals, stained for Discs large (Dlg),phospho-histoneH3, cleaved caspase 3 (inlet) and DNA (DAPI), Scale bars=50 *μ*m, 50 *μ*m (inlet). (**B**) Quantification of the number of 3^rd^ instar larval midgut AMP clusters and the number cells of within a given cluster of animals in (A).(**C**) Immunofluorescence of 3^nd^ instar larval midguts of *myo1A>, myo1A>ptcRNAi, myo1A>smoRNAi* end *esg>, esg>ptcRNAi, esg>smoRNAi* animals, stained for Discs large (Dlg), phospho-histoneH3, cleaved caspase 3 (inlet) and DNA (DAPI), Scale bars=50 *μ*m, 50 *μ*m (inlet). (**D**) Quantification of the number of 3^rd^ instar larval midgut AMP clusters and the number cells of within a given cluster of animals in (C). (**E**) Measurement of pupal volume of animals with increased or decreased Hh pathway activity in midgut ECs and AMPs. (**F**) Measurements of timing of pupariation of animals with increased or decrease Hh pathway activity in midgut ECs and AMPs (**G**) Survival rate upon starvation of *myo1A>ptcRNAi, myo1A>smoRNAi* and their respective control late 2^nd^ instar larvae.

**Supplemental Figure 4.**
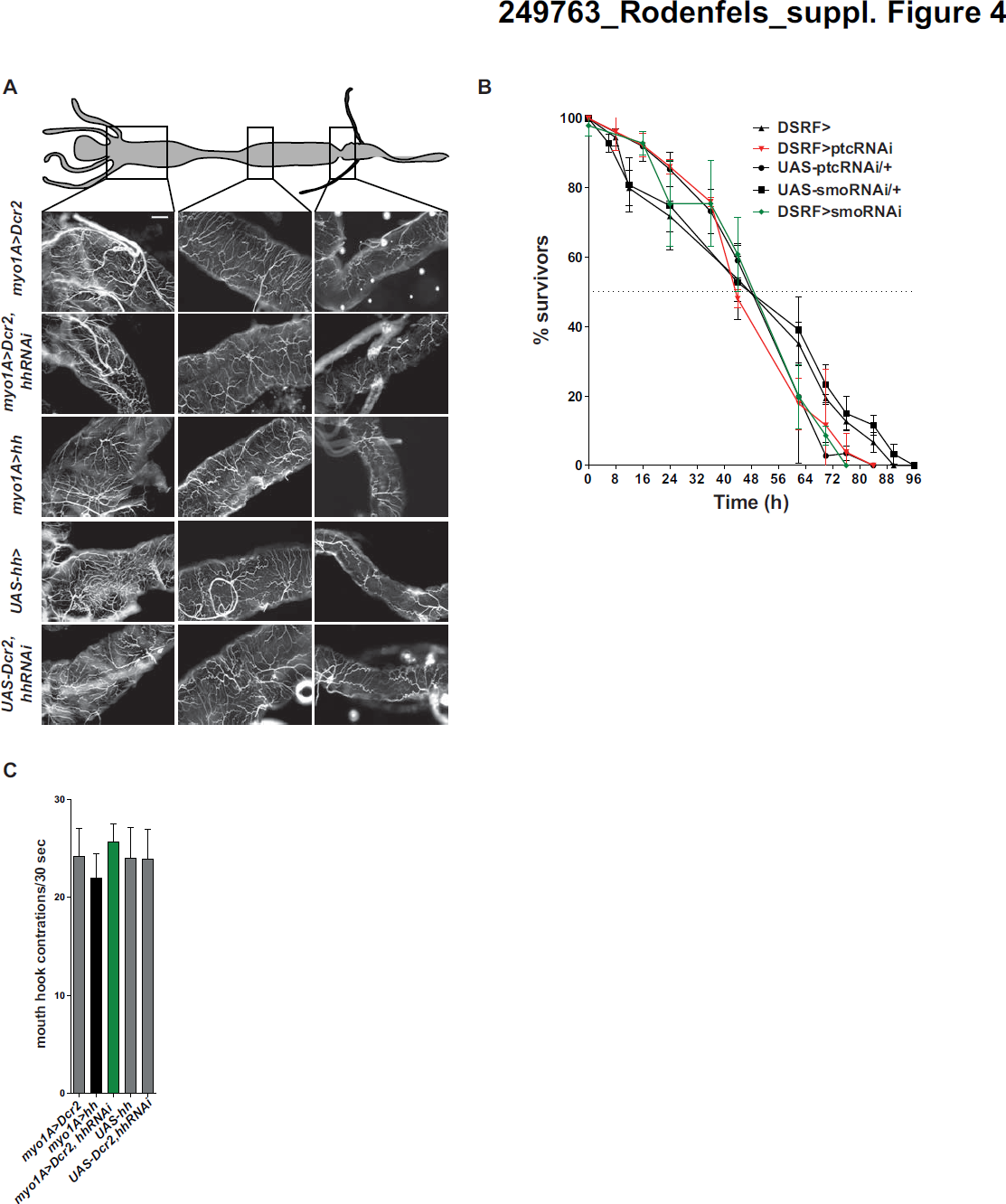
(**A**) Images of midgut trachea in 3 regions of early 3^rd^ instar midguts of 3 control genotypes (*myo1A>Dcr2, UAS-Dcr2;UAS-hhRNAi*, and *UAS-hh*) and *myo1A>Dcr2,hhRNAi* and *myo1A>hh* animals. (**B**) Survival rate upon starvation of *DSRF>ptcRNAi, DSRF>smoRNAi* and their respective control late 2^nd^ instar larvae (**C**) Quantification of the frequency of mouth-hook contraction of 3 control geneotypes (*myo1A>Dcr2, UAS-Dcr2; UAS-hhRNAi, UAS-hh*) and animals with increased (*myo1A>hh*) or decreased (*myo1A>Dcr2,HhRNAi) hh* expression in the midgut

**Supplemental Figure 5.**
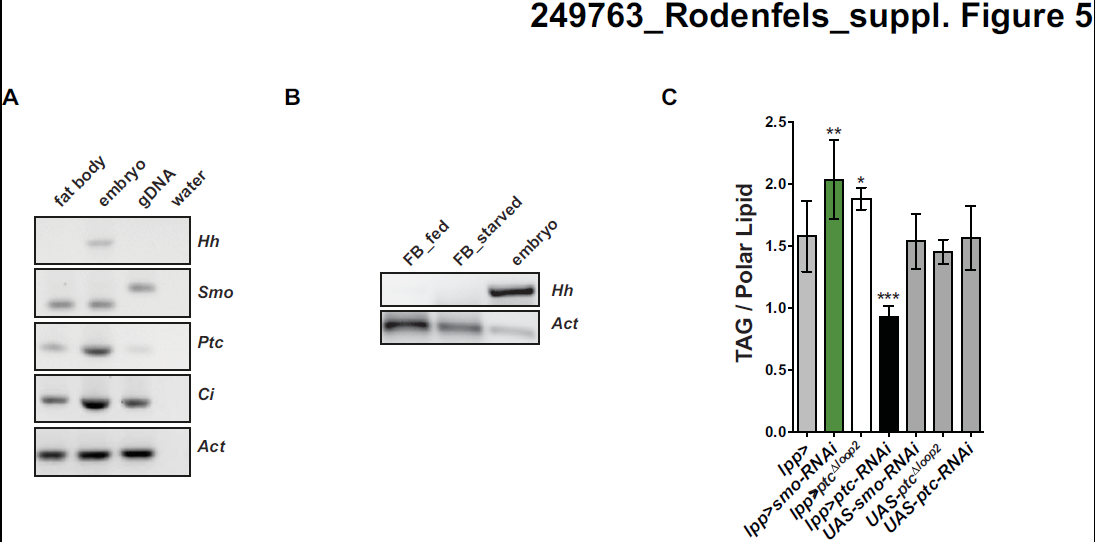
(**A**) RT-PCR for *hh, smo, ptc, ci* and *actin* from embryonic or 3^rd^ instar fat body samples. *Hh* message is absent whereas *smo,ptc* and *ci* meassages are present in fat body samples. (**B**) RT-PCR for *hh* from fed and starved fat body samples. *Hh* message is absent in fat body samples in every condition. (**C**) Whole animals TAG levels of normal fed late 3^rd^ instar (96h AEL) larvae with reduced or increased Hh pathway activity in the fat body quantified by mass spectrometry (n=3), normalized to total polar lipid. Increased Hh signaling mediated by the expression of *ptc*RNAi reduces TAG. Conversely decreased Hh signaling activity by *smo* knock-down or expression of *ptc*^Δ*loop2*^ causes increase in whole larval TAG.

**Supplemental Figure 6.**
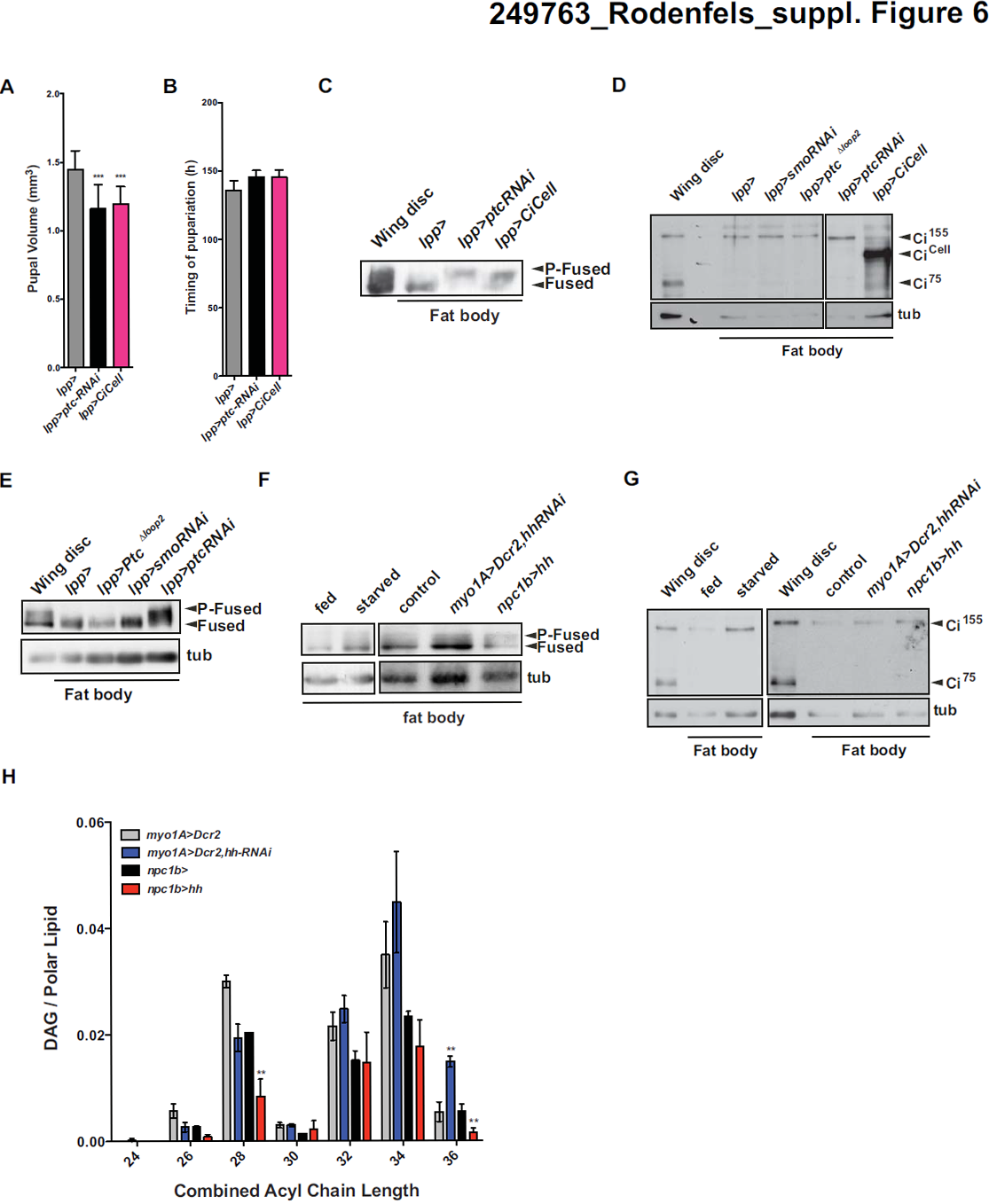
(**A**) Measurement of Pupal volume of *lpp>,lpp>ptc*RNAi and *lpp>CiCell* animals. (**B**) Measurements of timing of pupariation of *lpp>,lpp>ptc*RNAi and *lpp>CiCell* animals. (**C**) Immunoblot for wing disc or fat body Fused in 3^rd^ instar wild-type animals or larvae which express *ptc*RNAi or *CiCell* specifically in the fat body under the control of *lpp*–Gal4. (**D**) Immunoblot for wing disc or fat body Ci and tubulin (tub) in 3^rd^ instar wild-type animals or larvae which express *ptc*^Δ*loop2*^, *smo*RNAi, *ptc*RNAi and *CiCell* specifically in the fat body under the control of *lpp*–Gal4. (**E**) Immunoblot for wing disc or fat body Fused in 3^rd^ instar wild-type animals or larvae which express *ptc*^Δ*loop2*^, *smo*RNAi, *ptc*RNAi and *CiCell* specifically in the fat body under the control of *lpp*–Gal4. Tubulin was used as a loading control (**F**) Immunoblot for Fused of fed, starved and fed control, *myo1A>Dcr2,hhRNAi* and *npc1b>hh* fat body samples. Tubulin was used as a loading control. Neither condition changes the level of phosphorylated Fused. (**G**) Immunoblot for Ci and of fed, starved and fed control, *myo1A>Dcr2,hhRNAi* and *npc1b>hh* fat body samples. Tubulin was used as a loading control. Neither condition changes the ratio of Ci^155^ to Ci^75^. (**H**) Chain length distribution of fatty acid residues in DAG in fat body samples form fed 3^rd^ instar fat body samples from *myo1A>Dcr2, myo1A>Dcr2,hhRNAi, npc1b>* and *npc1b>hh* animals, quantified by mass spectrometry (n=3).

**Supplemental Figure 7.**
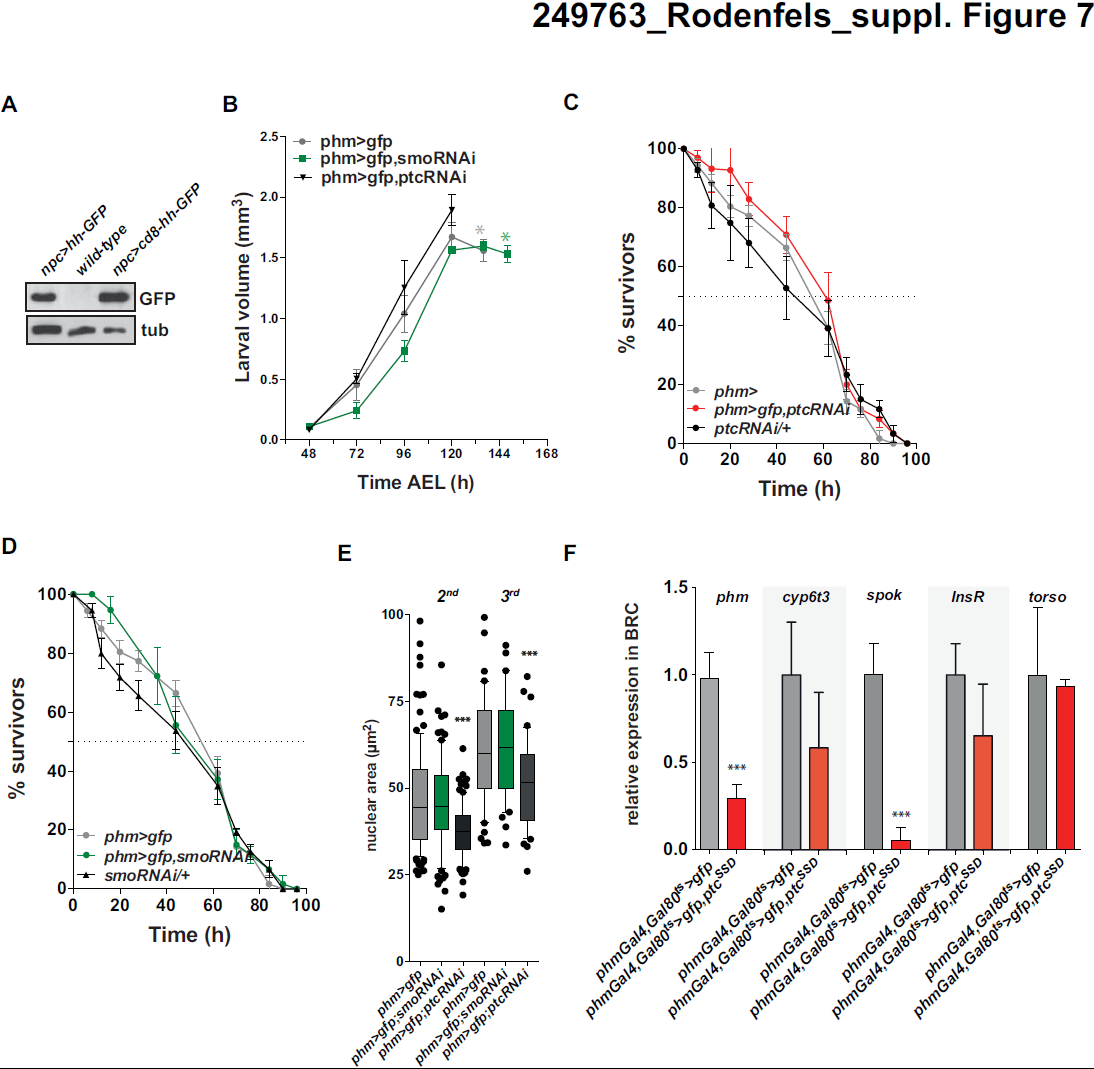
(**A**) Immunoblot for midgut GFP in 3^rd^ instar wild-type animals or larvae which express *hh-GFP* or cd8-hh-GFP specifically in the midgut under the control of *npc1b*– Gal4. (**B**) Changes in larval volume and determination of the timing of pupariation of *phm>gfp, phm>gfp,smoRNAi* and *phm>gfp,ptcRNAi* (A) and of animals with reduced (*myo1A>Dcr2,hhRNAi*) Asterix marks white pupal stage and volume (**C,D**) Survival rate upon starvation of *phm>gfp, phm>gfp,smoRNAi, phm>gfp,ptcRNAi, smoRNAi/+* and *ptcRNAi/+* late 2^nd^ instar larvae (**E**) Quantification of nuclear area of PG nuclei from phm>gfp, phm>gfp,smoRNAi and phm>gfp,ptcRNAi late 2^nd^ and 3^rd^ instar larvae, Error bars represent 95% confidence interval, ***p< 0.001 (**F**) Quantitative RT–PCR for *phm, cyp6t3, spok, InsR and toros* at 96h AEL from dissected brain-ring gland complexes with increased Hh pathway (*phmGal4,Gal80ts>gfp,ptc^SSD^*) activity in the PG relative to control (*phmGal4,Gal80^ts^>gfp*) BRC. Error bars represent SD of biological triplicates *p< 0.05, **p< 0.01 and ***p< 0.001.

## Supplemental Text

### Fat body Hedgehog Signaling Does not Depend on Ci Processing

We were surprised to find that fat body specific expression of CiCell, which constitutively mimics the repressive Ci^75^ and would be expected to increase pupal volume, instead has the same effect as pathway activation caused by knock-down of *ptc* – reducing pupal volume without affecting developmental timing (suppl. Figure 6A,B). *Ptc* transcription itself is normally activated by canonical Hh signaling. We therefore wondered whether the repressive CiCell might reduce growth indirectly, by lowering Ptc activity. If so, we would expect that *ciCell* expression should activate Hh pathway components lying downstream of Ptc but upstream of Ci processing. To test this idea we examined phosphorylation of Fused, a kinase required to regulate Ci processing that becomes phosphorylated upon activation of Smo (Therond et al. 1996; Ruel et al. 2007). Indeed, *CiCell* expression increases phosphorylation of Fused kinase, just like ptc-RNAi (suppl. Figure 6C). These observations confirm that *ciCell* expression activates more upstream components of the Hh pathway. This strongly suggests that Hh signaling in the fat body proceed via a non-canonical mechanism.

To investigate whether Ci processing is regulated by Hh signaling in the fat body, we first compared levels of Ci^155^ and Ci^75^ in wing discs and fat body. Wing discs accumulate approximately equal amounts of Ci^155^ and Ci^75^ (suppl. Figure 6D). In contrast, the Ci pool in the fat body consists almost entirely of Ci^155^. In the wing disc, stabilization of full length Ci^155^ depends on Smoothened signaling activity. Surprisingly, the extremely high ratio of unprocessed to processed Ci in the fat body does not, since neither smoRNAi nor expression of*ptc*^Δ*loop2*^ increases the amount of Ci^75^ (suppl. Figure 6D). Thus, Ci^155^ is constitutively stabilized in the fat body, independently of Hh signaling.

In the wing disc, Hh signaling increases phosphorylation of the serine-threonine kinase Fused, thereby blocking processing of Ci^155^ to Ci^75^. Fused is present in the fat body, and although it can be phosphorylated upon loss of Ptc, its phosphorylation under all other physiological conditions is barely detectable – much less than in the wing disc (suppl. Figure 6E). Thus, constitutive Ci^155^ stabilization in the fat body is not dependent on either Smo signaling or Fused phosphorylation.

Furthermore, although loss of Ptc does induce Fused phosphorylation in the fat body, different conditions that increase the amount of circulating Hh and induce physiological effects have no obvious effect on Fused or Ci (suppl. Figure 6F,G). Taken together, these observations indicate that circulating Hh signals to the fat body by a non-canonical mechanism that requires Ptc and Smo, but does not cause Fused phosphorylation or influences Ci processing. Recent data suggest that Smo can act as a G-protein coupled receptor (GPCR) (DeCamp et al. 2000; Ogden et al. 2008; Polizio et al. 2011a; Polizio et al. 2011b; Bijlsma et al. 2012). GPCR’s activate Ga and GB, Gy subunits to regulate second messengers such as cyclic AMP, Inositol–3–phosphate or diacylglycerol (DAG) (Wettschureck and Offermanns 2005). Interestingly, we find that the loss and gain of circulating Hh cause opposing effects on the levels of long chain DAG in the fat body (suppl. Figure 6H). This raises the possibility that non-canonical Smo signaling in the fat body may involve DAG as a second messenger.

## Supplemental Material & Methods

### Antibodies

Antibodies used: Hh (Eugster et al. 2007), Cv–D (Palm et al. 2012), Ci 1C2 (Wang and Holmgren 1999), Ci 2A1,Ptc, Smo, Fu, Dlg, tubulin (DSHB), phospho-HistoneH3, cleaved caspase 3 (Cell Signaling) and GFP (Invitrogen)

### Hemolymph Preparation

Diluted hemolymph was obtained by bleeding larvae with a tungsten needle in ice-cold PBS. Hemocytes were removed by centrifugation for 30 min at 1500g, 4°C. To remove cellular fragments, the supernatant was subsequently spun for 30 min at 16000g, 4°C. The supernatant from the second centrifugation step was snap–frozen in liquid nitrogen and stored at 80°C until use.

### Fly Husbandry and Fly Food

General methods for fly genetics and husbandry have been described (Ashburn et al. 2004). For all experiments flies were kept on standard cornmeal-yeast food or starvation media. Flies were kept under a 12 h light/12h dark cycle at 25 °C unless otherwise indicated. To deplete lipids from yeast autolysate, it was extracted with Chloroform 3 times overnight.

### Immunohistochemistry on Larval Tissues

Tissues were dissected form larvae at the indicated time AEL in 1× PBS on ice, fixed in 4% paraformaldehyde (w/v) in PBS for 20min at RT. Tissues were permeabilized with 0.05 % (v/v) Triton X-100 (PBX) for wing discs, 0.1 % PBX for ring glands and fat bodies and 0.2% PBX for midguts. Wing discs were blocked with 0.1 % (w/v) bovine serum albumin in PBX. Brain-ring gland complexes, fat bodies and midguts were block in 10 % (v/v) normal goat serum in PBX. Tissues were incubated with primary antibody over night in blocking solution at 4°C. After several washes in with the respective blocking solutions, tissues were incubated with secondary antibody and/or with rhodamin-phalloidin (Invitrogen) for 3–4h at RT. Wing disc were blocked additionally with 0.1% (w/v) and 4 % (v/v) normal goat serum for 30 min prior to the incubation with secondary antibody. Tissues were washed with PBX several times and incubated with DAPI for 5 min and mounted in either Prolong Gold Antifade reagent (Invitrogen) for wing discs or VectaShield (Vectalabs) for other tissues.

### Generation of UAS-hh-gfp and UAS-cd8-Hh-gfp

*UAS-hh-gfp* and *UAS-cd8-Hh-gfp* were generated using standard techniques. A 772 bp fragment coding HhN was cloned 5’prime of *gfp* cDNA. The resultant *hhGFP* cDNA fragment was fused at its 3’prime end either to a 4×MYC epitope (*hhGFP^MYC^*) or to a CD8 transmembrane domain sequence that is flanked at the 5’prime end with the 2×MYC-epitope and the 3’prime with a V5 epitope (*hhGFP^TM^*). Both constructs were cloned into a vector containing a 5×UAS-*hsp70* promoter element followed by a multiple cloning site, a SV40 element and mini-*white* gene. Transgenic fly lines were generated by phiC31-mediated integration of UAS-*hh*GFP^*MYC*^ and *hh*GFP*^TM^* into VK33 *attP* landing site (Markstein et al. 2008).

### *In-Situ* Hybridization

*In-situ* hybridization with digoxigenin-labeled probes was essentially performed as previously described (Tomancak et al. 2003). Briefly, DNA templates were generated by PCR and purified. The purified PCR product was transcribed by incubation for 2h at 37°C after the addition of 5 μl of a polymerase reaction cocktail consisting of 2 U T7 RNA polymerase, 4.6 U RNase inhibitor, 10 mM NTPs, 3.5 mM digoxigenin-11-UTP, 40 mM Tris pH 8.0, 6 mM MgCl_2_, 10 mM DTT and 2 mM spermidine. After treatment with DNase I and Na_2_CO_3_ pH 10.2, ethanol precipitation was carried out. Pellets were resuspended in 50 μl of 50% formamide, 5 mM Tris-HCl pH 7.5, 0.5 mM EDTA and 0.01% Tween 20. Re-hydrated and fixed midguts were incubated for 1 h in hybridization buffer (50% formamide, 4× SSC, and 0.01% Tween 20). Digoxigenin-labeled RNA probe (1:100 dilution in hybridization buffer with 5% dextran sulfate) was added and the midguts were incubated overnight at 55°C. The midguts were subjected to eight 30-min washes in wash buffer (50% formamide, 2x SSC and 0.01% Tween 20). Midguts were treated for 2 h with 5% goat serum and anti-digoxigenin-AP Fab fragments (Roche). Following nine 10-min washes in 0.1% Tween 20 in PBS and two rinses in AP buffer (50 mM MgCl_2_, 100 mM NaCI, 100 mM Tris pH 9.5, 0.01% Tween 20) the NBT/BCIP color substrates were used to detect the hybridized probes. Midguts were washed six times with ethanol to enhance contrast and stored in 70% glycerol in PBS.

### RNA extraction and reverse transcription polymerase chain reaction

Larval fat bodies or midguts were dissected in ice-cold PBS and frozen in liquid nitrogen. Total RNA was extracted using a Qiagen RNeasy lipid tissue kit according to manufacturers protocol. RNA samples were treated with DNase I for 40 min at 37° C and purified again with a Qiagen RNeasy kit according to manufacturers protocol. RNA samples were reverse transcribed using SuperScript Vilo cDNA synthesis kit (Invitrogen). PCRs were performed according to standard protocols with equivalent amounts of cDNA from each tissue.

### Quantitive RT-PCR

Generated cDNA samples were used for real-time RT-PCR using SYBR-Green PCR Mastermix (Thermo Fisher Scientific) with 2.5 ng of cDNA template and efficiency optimized primer were used at concentrations between 100 nM-300 nM. Samples were normalized to *rp49*. Six independent samples were collected for each experiment and triplicate measurements were conducted. Primer sequences are available upon request.

### Quantification of Mouth-Hook Contractions

The quantification of the frequency of early 3^rd^ instar larval mouth-hook contractions on solid foods was performed as described (Wu et al. 2005).

### Imaging of Midgut Trachea

3^rd^ instar midguts were dissected and immediately mounted in 100% glycerol and imaged by dark field microscopy. Trachea appear as bright branched structures covering the midgut.

### Quantification of Midgut Adult Midgut Progenitor Cell (AMP) Clusters

Midgut AMPs and clusters, which lie on the basal side of the gut epithelium, are diploid and their nuclei differ in size from those of polyploidy enterocytes and endoendocrine cells (Jiang and Edgar 2009). This allows identification on the basis of by nuclear DNA staining. The number of AMP clusters and cells within a given cluster were counted from a particular region of interested of the posterior midgut for n>10 midguts for a particular genetic background.

### Statistical analysis

Raw data was initially processed using Microsoft Excel. Plotting and statistics of the processed data sets were performed using GraphPad Prism version 5.0c for Mac OS X, GraphPad Software, San Diego California USA. Mathematical significance of differences between datasets was analyzed using the unpaired t test and expressed as p-values.

